# The genetic sequences prone to copy number variation and single nucleotide polymorphism are linked to the repair of the poisoned DNA topoisomerase II

**DOI:** 10.1101/2020.09.03.280669

**Authors:** Chuo Jiang, Cong Ma, Detao Wang, Li Liu, Chunxiu Zhang, Fuxue Chen, Jiaxi Wu

## Abstract

TOP2-poisoning bioflavonoids and pesticides are linked to the copy number variation-related autism and chromosome translocation-related leukemia. On the other hand, the poisoned DNA topoisomerase II (TOP2) can lead to chromosome aberration. However, except a limited number of genes such as the *MLL* fusion, other poisoned TOP2-targeted genes, as well as their relationships with any specific diseases, are not defined. We applied the γH2A.X antibodies to genome-widely immunoprecipitate the chromatins that were associated with the repair of the TOP2 poison etoposide-induced DNA double strand breaks. We identified many transcriptable protein- and nonprotein-coding DNA sequences that are the candidates of or associated with many gene copy number variation- and/or single nucleotide polymorphism-associated diseases, including but not limited to microdeletion and microduplication syndromes (which are phenotypically presented as developmental, autistic, neurological, psychiatric, diabetic, autoimmune, and neoplastic diseases among many others) as well as stature, obesity, metabolic syndrome, hypertension, coronary artery disease, ischemic stroke, aortic aneurysm and dissection, leukemia, cancer, osteoporosis, Alzheimer disease, Parkinson disease, and Huntington disease. Our data raise the possibility that the poisoned TOP2 might be linked to the specific genetic alterations contributing to these diseases, additional to the known copy number variation-related autism and chromosome translocation-related leukemia. According to our and others’ data, we propose a model that may interpret the features, such as mosaicism, polygenic traits and pleiotropy, of these diseases.

**Author Summary:** For the past several decades, the morbidity rate of many diseases, including autism, mental disorders, cancer, cardiovascular diseases, diabetes, and senile dementia, has world-widely been rising. Analysis of the genome of the patients and their family members has identified the genes, whose alterations, so called copy number variation (CNV) and single nucleotide polymorphism (SNP), contribute to the diseases. Moreover, the CNVs and SNPs are *de novo*, that is, they have occurred only in the recent generations. Epidemiologically, this indicates that for the past several decades, there have existed some unknown world-wide etiologies to which human beings are exposed. If the etiologies are identified, avoiding human’s exposure may reduce the morbidity of the diseases. We have found that the repair of the poisoned topoisomerase II involves many genes that contribute to the aforementioned diseases. As the topoisomerase II is known to be located at the genomic sites where the disease-associated CNVs occur, as the poisoned topoisomerase II is susceptible to chromosome aberration, and as the topoisomerase II poisons, such as dietary bioflavonoids, are widely distributed in the environment, our data raise the yet-to-be-confirmed possibility that the environmental topoisomerase II poisons might etiologically contribute to many CNV-associated diseases.

## Introduction

γH2A.X is a DNA repair-involved, post-translationally modified form of histone variant H2A.X [1–4]. When a DNA double strand break (DSB) is elicited, H2A.X is phosphorylated on its serine 139 to generate γH2A.X as an early cellular DNA repair response [1–4]. γH2A.X molecules can spread up to megabases along the DNA flanking the DBS [1–4]. Previous studies have shown that the endogenous DSB causes the *in situ* formation of γH2A.X from the H2A.X that is located around the sub-telomeric region and transcription starting site in the Jurkat cell [5]. On the other hand, the exogenous DSB induced by 10Gy ionizing irradiation leads to the distribution of γH2A.X at the pausing site of RNA polymerase II within the gene body in the resting normal CD4 cell [5].

Human DNA topoisomerase II (TOP2) is an enzyme that relieves the topological strain generated during replication, transcription and other DNA transactions [6–8]. The enzyme may be poisoned by both distorted DNA structures, such as abasic DNA and DNA adducts, and many endogenous and exogenous agents, such as reactive oxygen species and bioflavonoids [6–8]. The poisoned TOP2 is converted by the cell into a TOP2-linked DSB known as cleavable complex [6–8], which subsequently triggers a series of cellular DNA repair responses, including the formation of γH2A.X [3,9,10].

Additional to the DSB, the poisoned TOP2 may also cause a DNA single strand break (SSB) [11]. However, the SSB does not seem to be related to the formation of γH2A.X and cytotoxicity [12]. More importantly, the poisoned TOP2-idnuced chromosome aberration, on which the current study focuses, is considered to result from the DSB, not the SSB [13].

TOP2 poisons may cause chromosome aberration [6–8,14,15]. Additionally, the TOP2-poisoning anticancer drug etoposide, bioflavonoids and pesticides have been linked to the chromosome translocation-related leukemia and/or gene copy number variation (CNV)-related autism [7,13,16–18]. However, except a limited number of genes such as the leukemia-related *MLL* fusion, other mutated genes resulting from the TOP2 poison-induced DNA damage and, particularly its repair, are not defined. Nor are their relationships with any specific diseases.

Mechanistically, the formation of the *MLL* fusion is considered to be caused by the DSB induced by the poisoned TOP2 [13], apoptotic caspase-activated DNase (CAD) [19,20] and/or other mechanisms [21,22]. However, clinically, patients who have the *MLL* fusion and develop leukemia, often have a history of receiving the TOP2-poisoning anticancer drugs [23].

Additional to the autism, CNVs are attributed to a group of diseases named as microdeletion and microduplication syndromes (or genomic disorders) which are characterized by the deletion or duplication of a genomic segment that often encompasses a number of transcriptable protein- or nonprotein-coding sequences [24–28]. (Hereafter, the terms of “transcriptable protein- or nonprotein-coding sequences”, “transcriptable sequences”, “genetic sequences”, and “genes” will be used interchangeably.) These syndromes often have complexed phenotypes and are presented as developmental, autistic, neurological, psychiatric, diabetic, autoimmune, and/or neoplastic diseases [24–28].

To identify the genes that are related to the repair of the poisoned TOP2, as well as their relationships with any specific diseases, we applied the γH2A.X antibody-dependent chromatin immunoprecipitation-DNA sequencing (ChIP-seq) technology to genome-widely capture Jurkat cell’s genetic sequences that were associated with the repair of the etoposide-induced DSBs. We determined to precipitate the chromatins associated with the DNA repair rather than those only associated with the TOP2 cleavable complexes, for the reason that the poisoned TOP2-induced chromosome aberration often results from the DNA repair that involves DNA sequences far from the TOP2 cleavable complexes [8,29]. As expected, we captured many genetic sequences that are known to contribute to leukemia and autism. However, unexpectedly, a large number of genetic sequences that are related to many other diseases, were also identified. The possible roles of the poisoned TOP2 in the pathogenesis of these diseases are discussed and a hypothetical model is proposed to interpret the complexed features of these diseases.

## Results

Poisoned TOP2 has long been known to cause chromosome aberration. However, the vast majority of the involved genes are undefined. Therefore, the only purpose of the current study was to identify the genes that were associated with the repair of the poisoned TOP2 and to find if these genes were related to any specific diseases. It was not our intention to study the chemistry, biochemistry or biology of the poisoned TOP2 or its targeted genes, because a large number of pioneering investigators have done many explicit studies. As a result, we only present the data that are related to the genetic sequences and the associated diseases. We do not show those sophisticated and exquisite figures or data that are generated by the ChIP-seq computer software, the usual way that most ChIP-seq studies do. We would not like to distract the readers from the main theme of the current study.

### ChIP captured a number of transcriptable protein- and nonprotein-coding genetic sequences

We treated the human leukemia Jurkat cells continuously with 100μM etoposide, waited for hours and then immunoprecipitated the chromatins with the anti-γH2A.X antibodies. We also immunoprecipitated the chromatins of the cells that were mock-treated with DMSO, which was used to prepare the stock solution of etoposide, to control any endogenous DSBs such as those described by Seo *et al* [5]. We designed our experiments in such ways for the following reasons.

1. Unlike other TOP2 poisons such as the bioflavonoids that have multiple cellular effects, including antioxidant, pro-oxidant, anti-mutagenic, mutagenic, carcinogenic, and anti-carcinogenic activities, which have not been thoroughly characterized at the molecular level [13], etoposide is well known to poison TOP2 and hence has been used by most investigators to study the poisoned TOP2 [6–8]. We utilized etoposide just as a TOP2-poisoning tool because our only purpose was to identify the genetic sequences associated with the poisoned TOP2 and its repair. We did not intend to characterize any specific TOP2 poisons. Therefore, the principle of the conclusion of the current study should be applicable to other TOP2 poisons such the TOP2-poisoing bioflavonoids, although the details are not likely to be exactly the same because of the disparate efficacy, potency and DNA sequence selectivity among various TOP2 poisons.
2. Early studies have immunoprecipitated the TOP2 cleavable complex- or DNA damage-associated chromatins [31,32]. We instead immunoprecipitated the γH2A.X- or DNA repair-associated chromatins, because the chromosome aberration induced by TOP2 poisons often occur as a result of DNA repair and involves DNA sequences far distant to the DSB [8,29]. Actually, the chromosome aberration is a direct result of the repair (such as homologous recombination and non-homologous end joining) or mis-repair of the poisoned TOP2-induced DSB, rather than the DSB itself. More importantly, the poisoned TOP2 in the TOP2 cleavable complex is quickly degraded through both ubiquitin-dependent and independent proteasome pathways [33–35]. Moreover, γH2A.X, which only binds to the naked DNA [1–4], is not like to bind to the DNA that is still occupied by the intact or partially degraded TOP2. Therefore, any ChIP-seq studies using the anti-TOP2 antibodies to precipitate the TOP2 cleavable complexes, are likely to underestimate the number of the TOP2 cleavage sites as well as the DNA repair-involved genetic sequences. Indeed, as shown by Dellino *et al.*, the regions enriched with the TOP2B ChIP-seq signals upon the etoposide treatment, do not correlate with those of the γH2A.X ChIP-seq signals [32].
3. Additional to the DSB, etoposide has also been shown to cause the poisoned TOP2-linked SSB [11,12]. However, we only addressed the TOP2-linked DSB, rather than the SSB, because it is the DSB that is considered to be associated with the TOP2 poison-induced chromosome aberration [13], γH2A.X formation and cytotoxicity [12]. Moreover, studying the chemistry, biochemistry or biology of the poisoned TOP2 was not the purpose of the current study.
4. We chose γH2A.X as an indicator of DNA repair, because it is formed at the early stage of DSB repair [1–4]. This way, we avoided the complexed, distinctive cellular outcomes (such as apoptosis, sentences, survival with or without chromosome aberrations, and differentiation). These distinctive cellular outcomes may result from different amounts of DNA damage and/or disparate DNA repair capabilities (i.e. whether the DSB is unrepaired, error-free repaired or error-prone repaired).
5. To capture as many genomic segments that were undergoing repair as possible, we employed 100μM etoposide and waited for 5.5 hours before collecting the cells for ChIP-seq. We treated the cells with 100μM etoposide because we concerned that a low dose of etoposide might not induce the sufficient amount of a signal that fell into the sensitivity range of the ChIP-seq method [5]; the amount of the signal of any single gene among a heterogenous cell population is likely to be low. We harvested the cells after the 5.5-hour drug treatment because our kinetic analysis showed that γH2A.X peaked at around 5 hours following the 100μM etoposide treatment (Supporting Information S1 Fig). It should be noted that the etoposide-induced *MLL* fusion seems independent of the dosage of etoposide as a wide range of doses, from 0.14-0.5μM, of etoposide can induce the fusion [36,37]. It should also be noted that DNA repair is a kinetically and dynamically changing process [1–4]. Therefore, the γH2A.X-associated DNA is not likely to be the same throughout the entire DNA repair process. Therefore, the genetic sequences captured by our ChIP only reflected those that were associated with γH2A.X at 5.5 hour after the etoposide treatment. On the other hand, as we discuss in the following Discussion section, these experimental conditions do not negate our conclusion.

Under the above discussed experimental conditions, a total number of 3707 transcriptable protein- and nonprotein-coding sequences were reproducibly or repeatedly identified by the ChIP-seq (NCBI SRA accession number SRP150381 at https://www.ncbi.nlm.nih.gov/sra).

Initially, we investigated these 3707 genetic sequences one by one to find whether they were related to any specific diseases. Three databases, Gene (https://www.ncbi.nim.nih.gov/gene/), PubMed (https://www.ncbi.nim.nih.gov/pubmed/) and Database of Genomic Variants (http://dgv.tcag.ca/dgv/app/home) were searched. We spent more than 6 months only to screen the first 500 or so chromatin-immunoprecipitated (ChIPed) genetic sequences. We found that at least 50% of them were linked or susceptibly linked to certain known diseases. However, this initial gene-disease correlation study was very laborious and quite inefficient. Additionally, none of the three databases seemed inclusive or updated. More importantly, for a large portion of our ChIPed genetic sequences, scientists have not seemed to reach a consensus on their gene-disease relationships. As a result, we changed our strategies. Instead of investigating every ChIPed genetic sequence, we spent another 6 months or more to focus on the genes of 17 diseases or conditions. The gene-disease relationships of these genes had well been documented in the literature. We chose these 17 diseases also for the reason that since the industrialization, most CNVs and SNPs related to these diseases have occurred *de novo* and the morbidity rates of these diseases have continuously been rising. We contemplated that preventive measures can be introduced to reduce the morbidity rates if the contributing etiologies to the genetic alterations of these diseases are identified.

### The poisoned TOP2-induced, γH2A.X-associated DNA was closely linked to many genetic sequences associated with the CNV-related microdeletion and microduplication syndromes

Microdeletion and microduplication syndromes refer to a group of diseases that arise from the deletion, duplication or other types of chromosome aberrations of a genomic segment, often leading to CNVs. Clinically, these syndromes may be manifested as developmental, autistic, renal, cardiovascular, neurological, psychiatric, autoimmune, neoplastic, and/or diabetic diseases [24–28]. Our ChIP captured many genetic sequences known to be within the deleted or duplicated chromosome segments of the syndromes. Table 1 lists these CNV-associated syndromes, as well as the involved genes, including their chromosome loci, which were captured by our ChIP. (The genes listed are not necessarily the candidate genes of the diseases and we only list one involved gene although, many times, more genes in the deleted or duplicated genomic segments were pulled down by the ChIP.) As illustrated, the genetic sequences of 66.4% (156 out of 235) of the microdeletion and microduplication syndromes reported by Weise *et al.*, were captured by our ChIP (Table 1) [27]. Therefore, the poisoned TOP2-induced, γH2A.X-associated DNA was closely linked to many CNV-related microdeletion and microduplication syndromes.

**Table 1.**
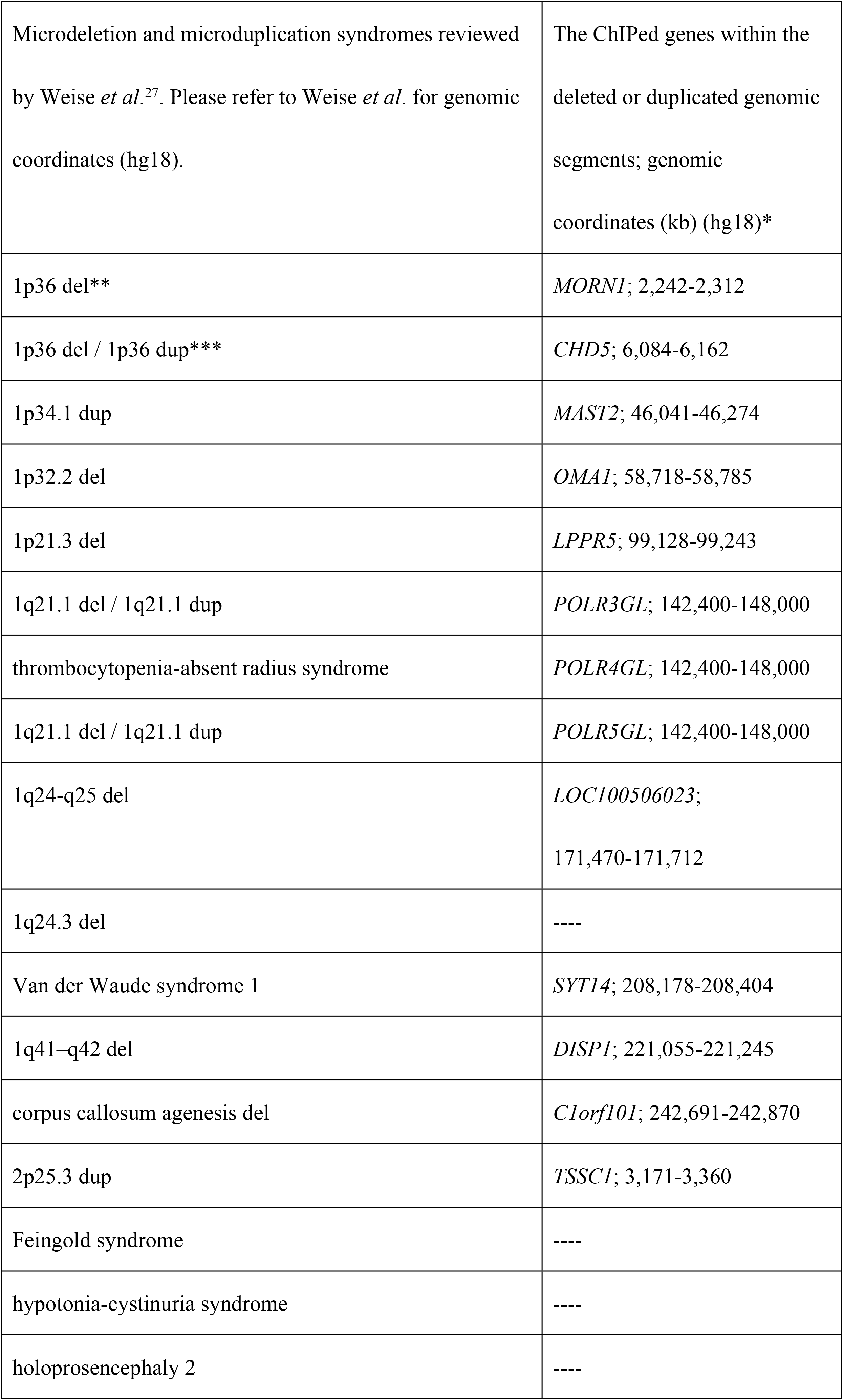
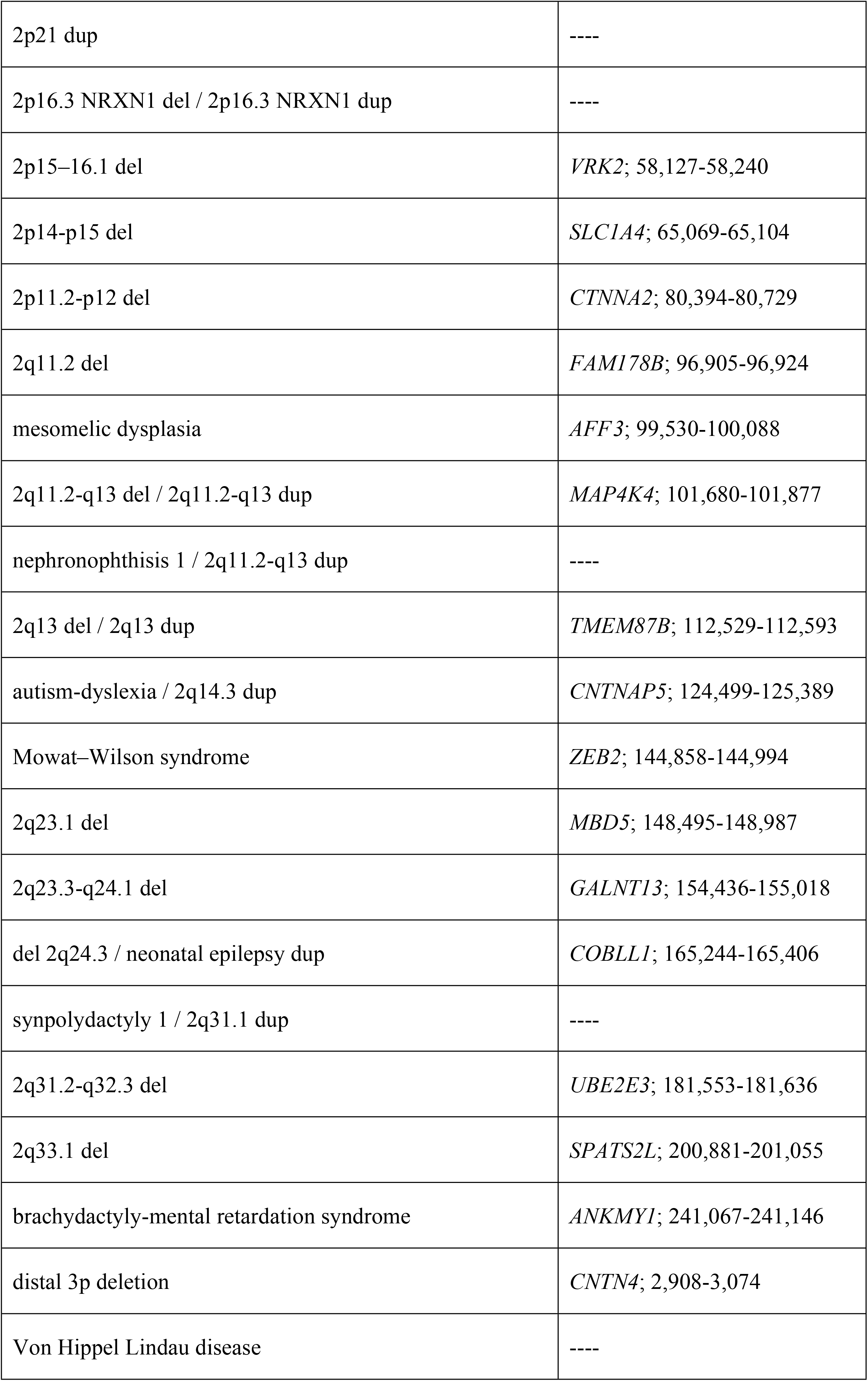
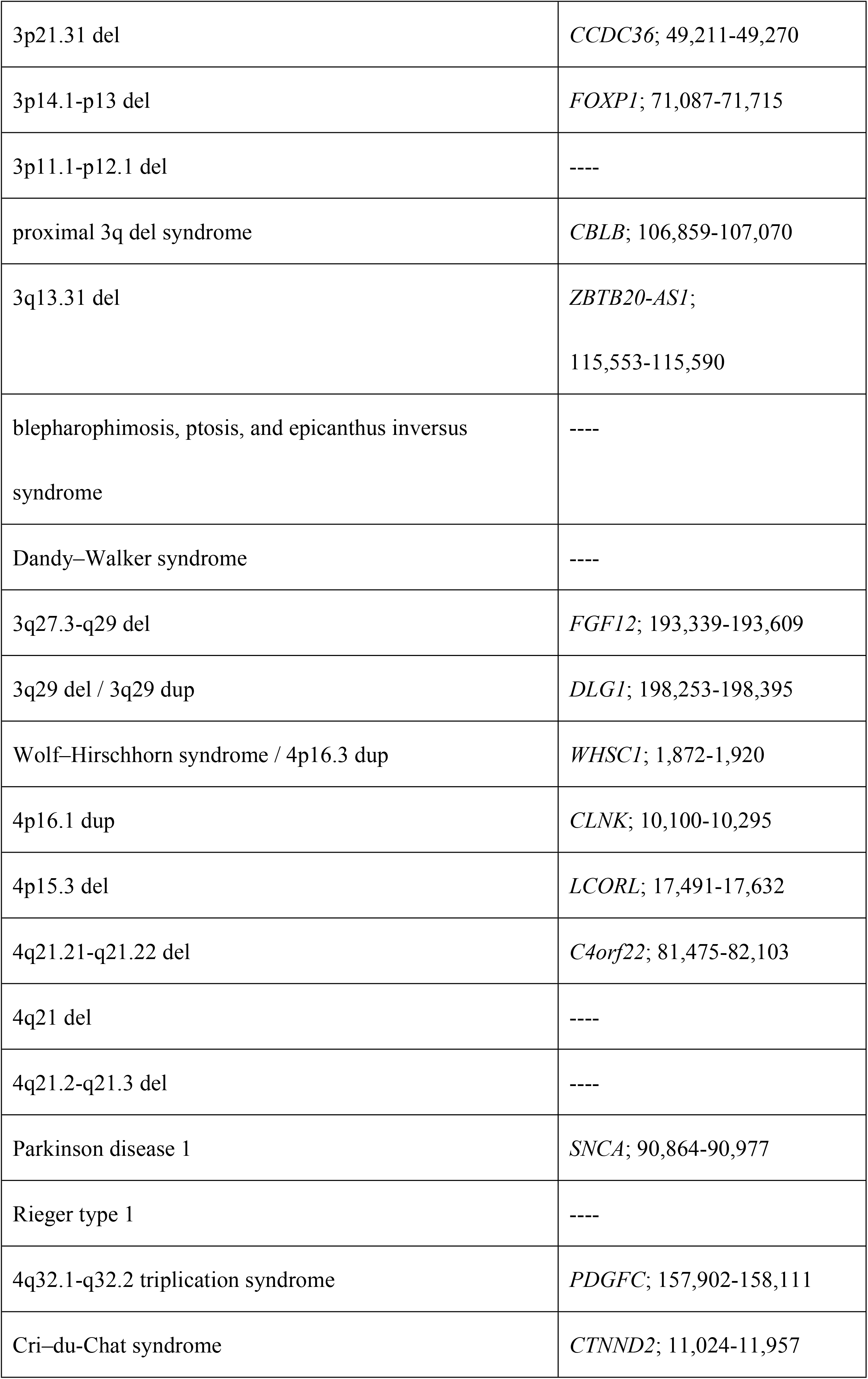
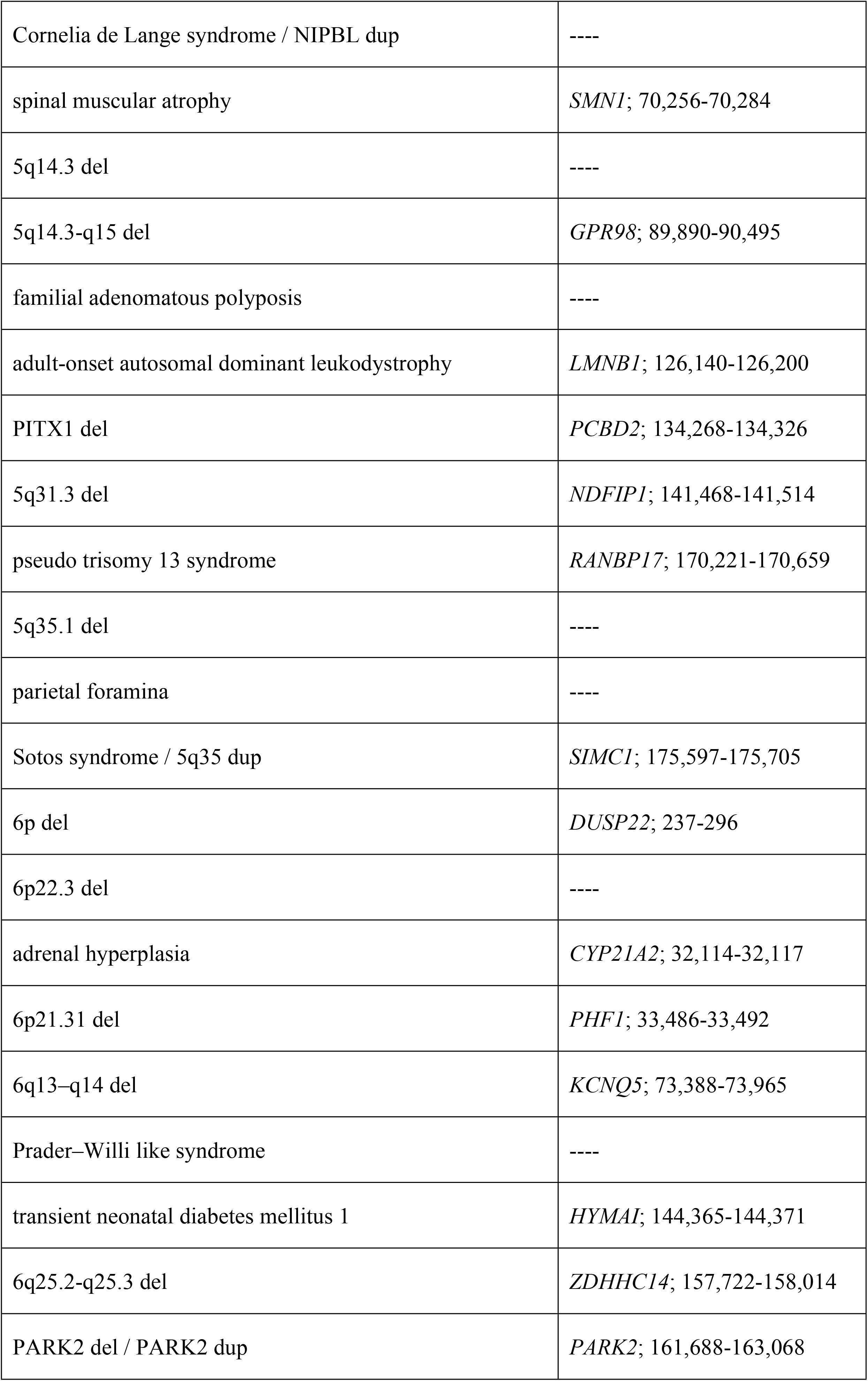
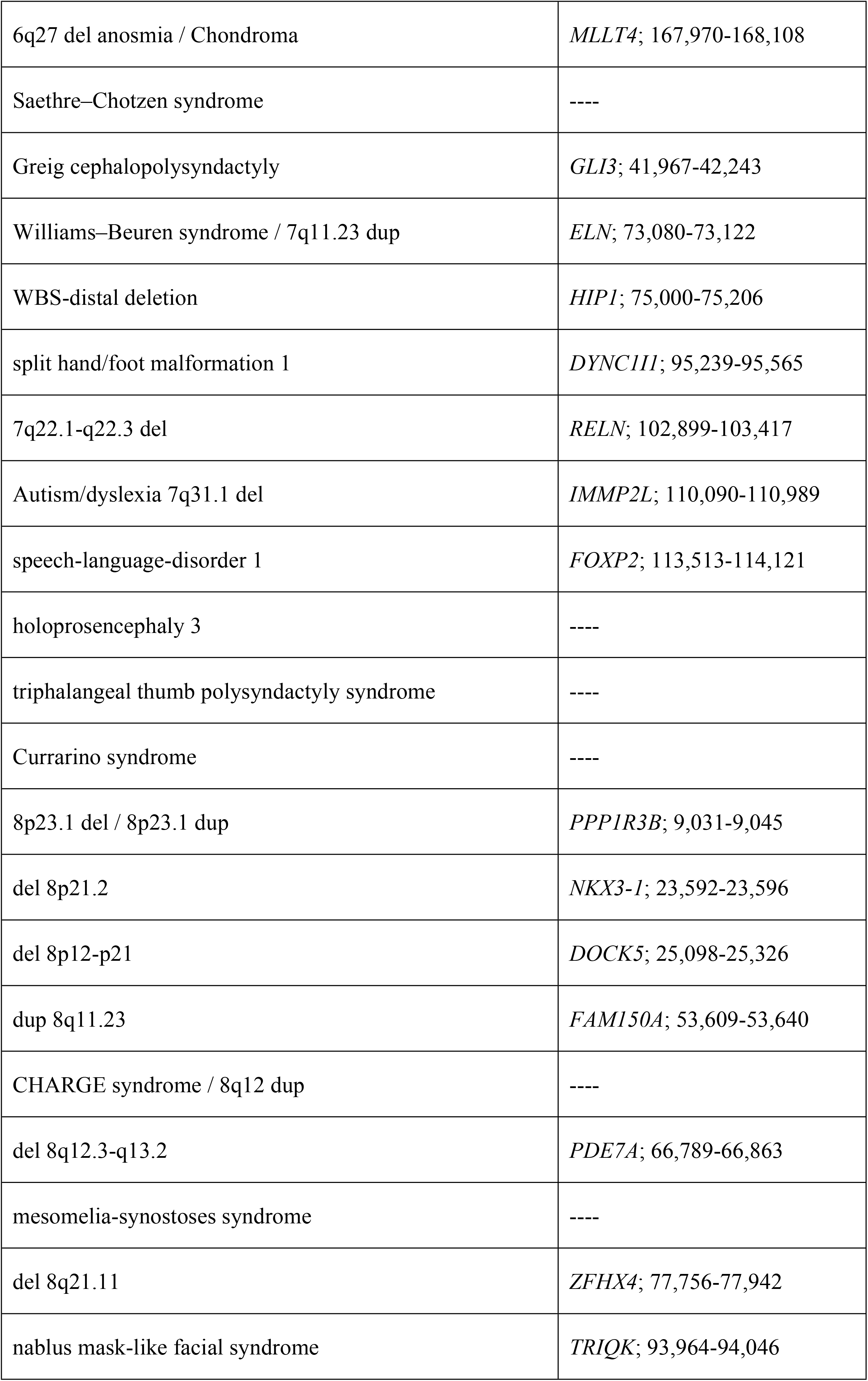
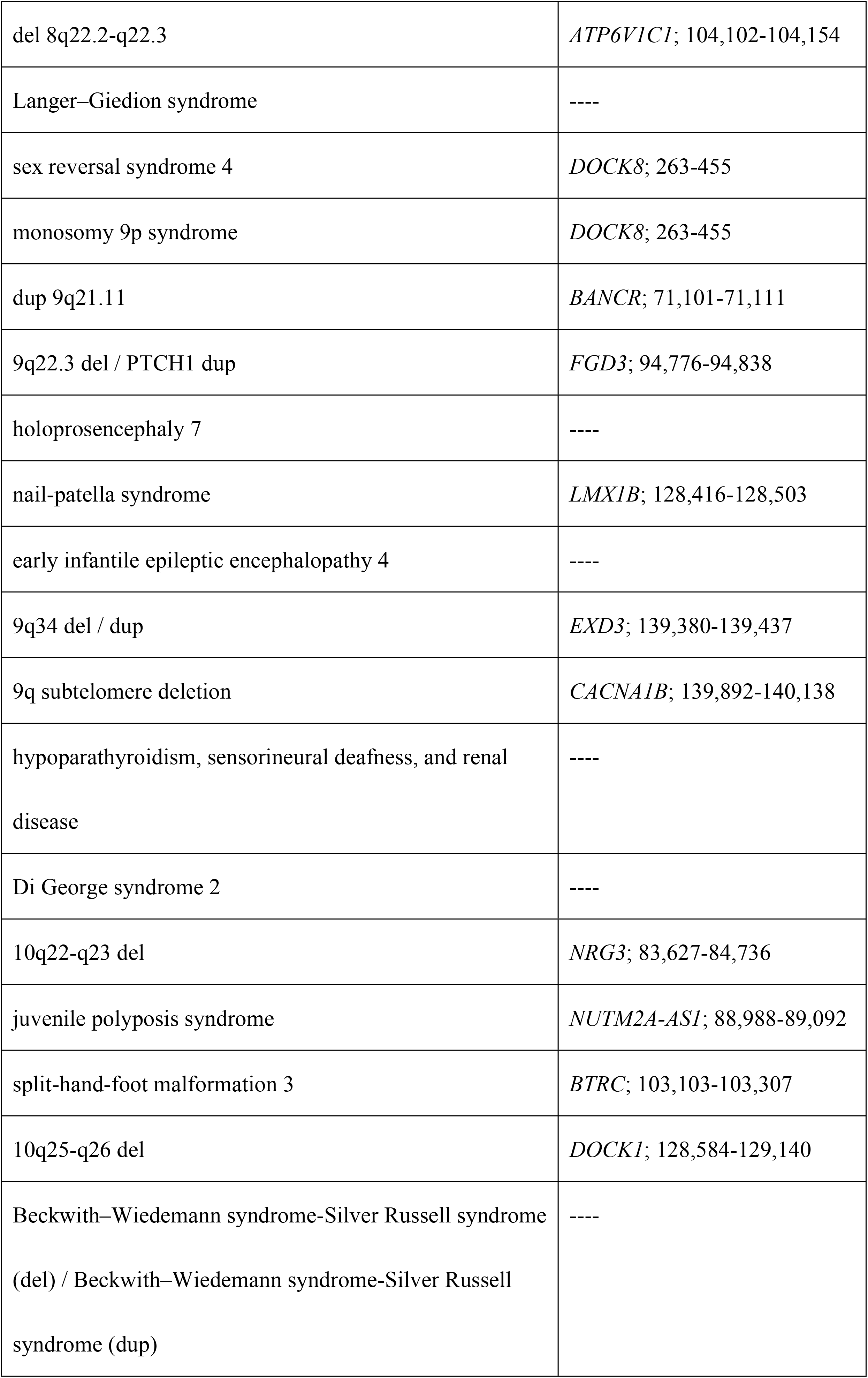
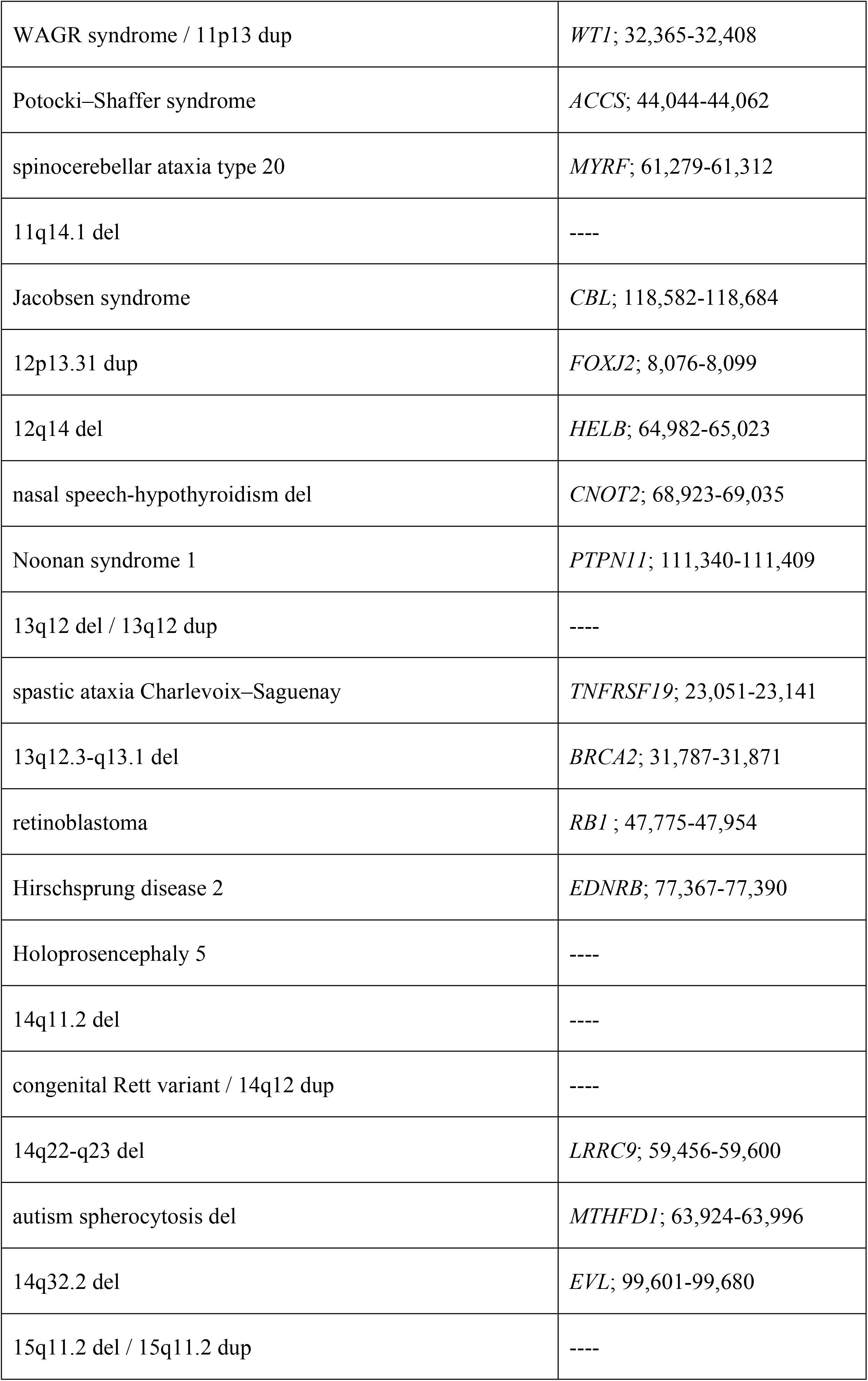
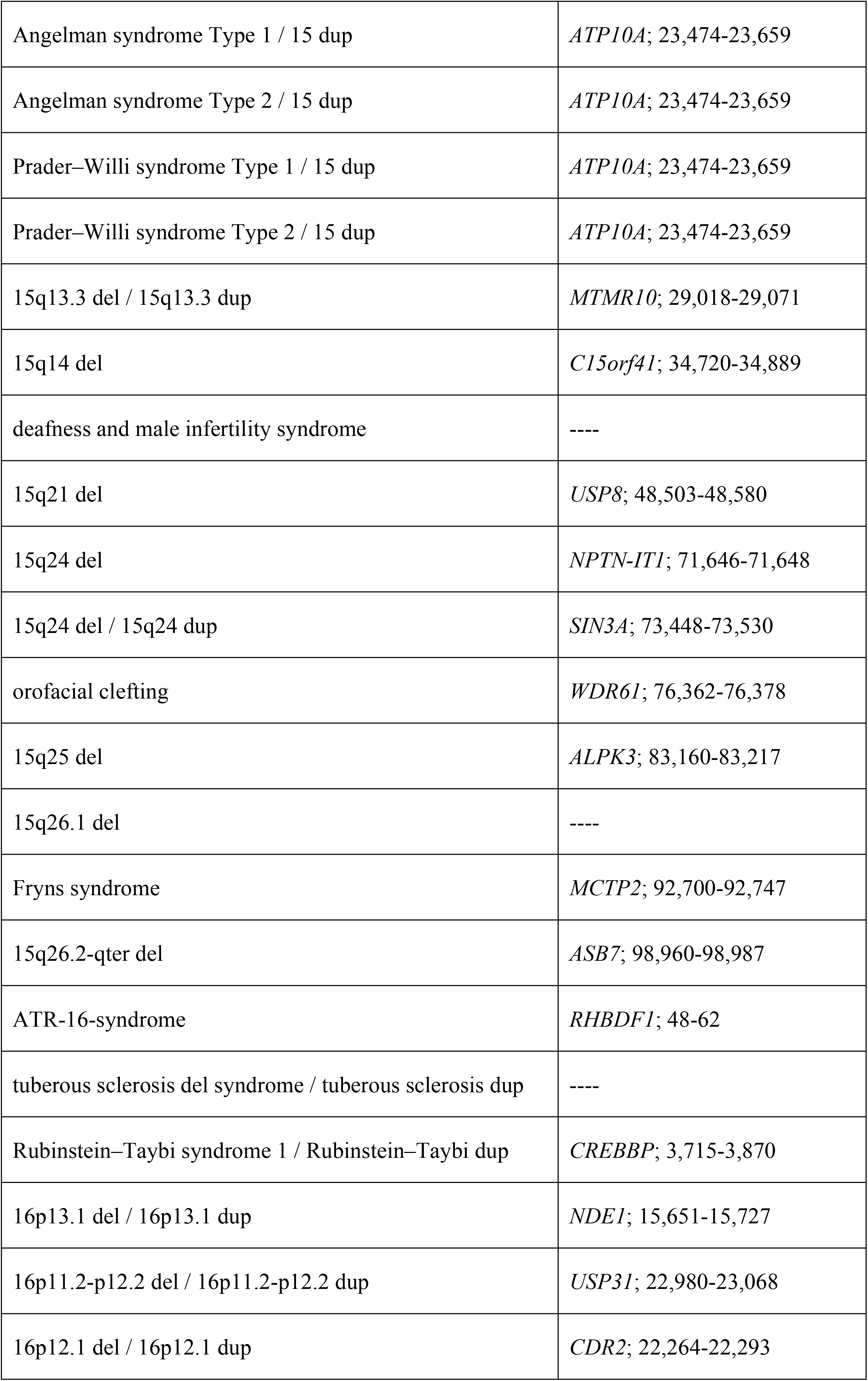
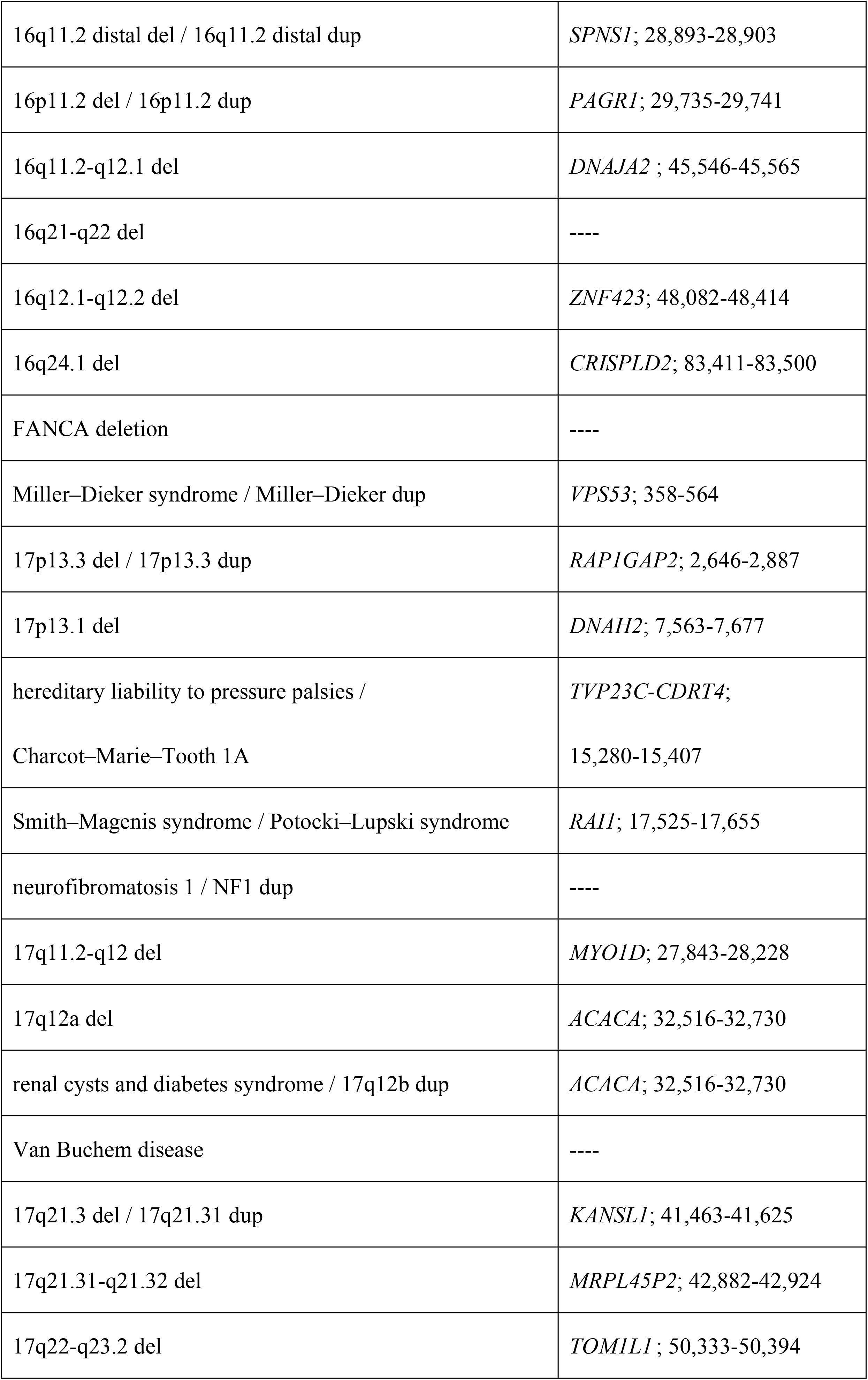
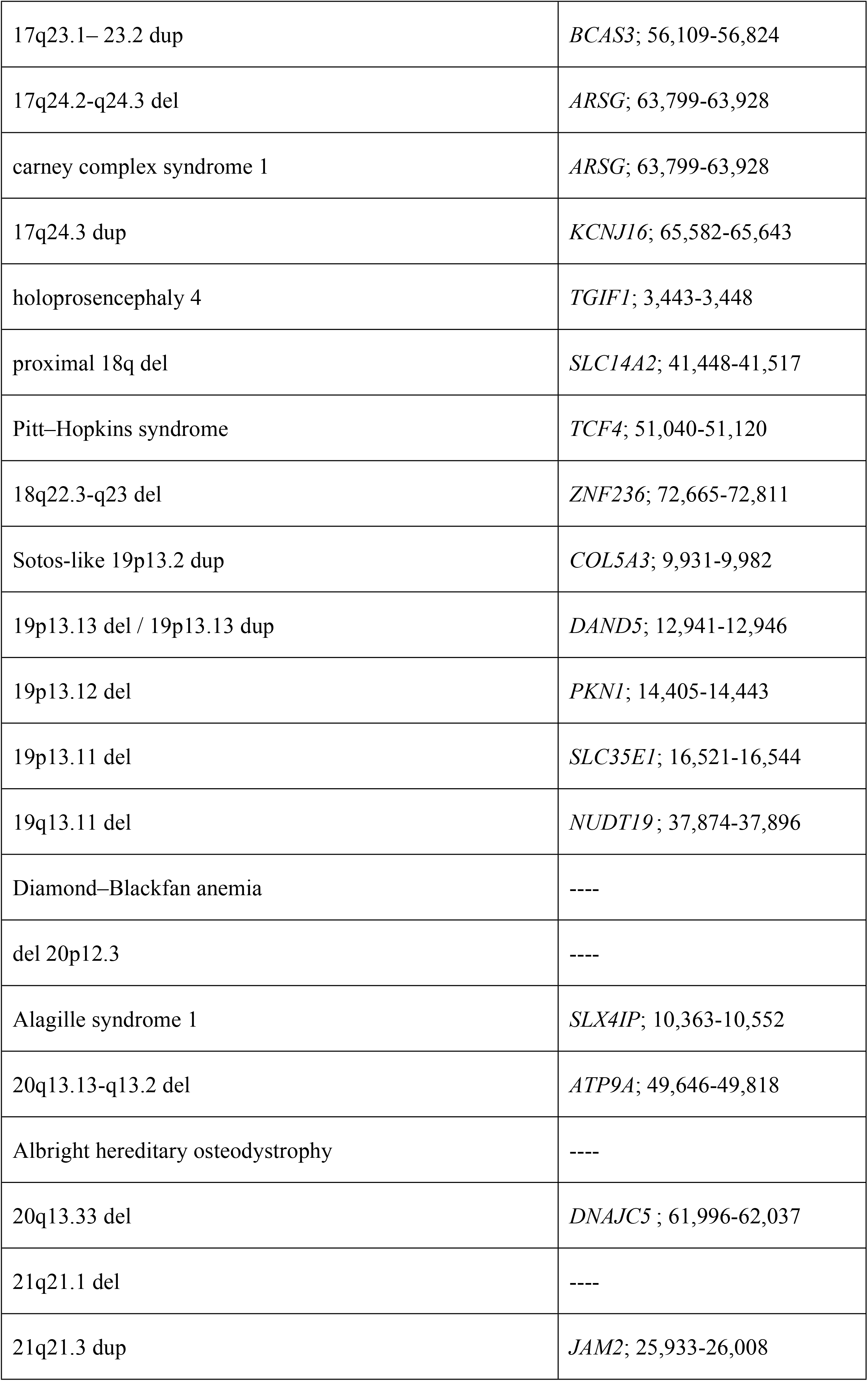
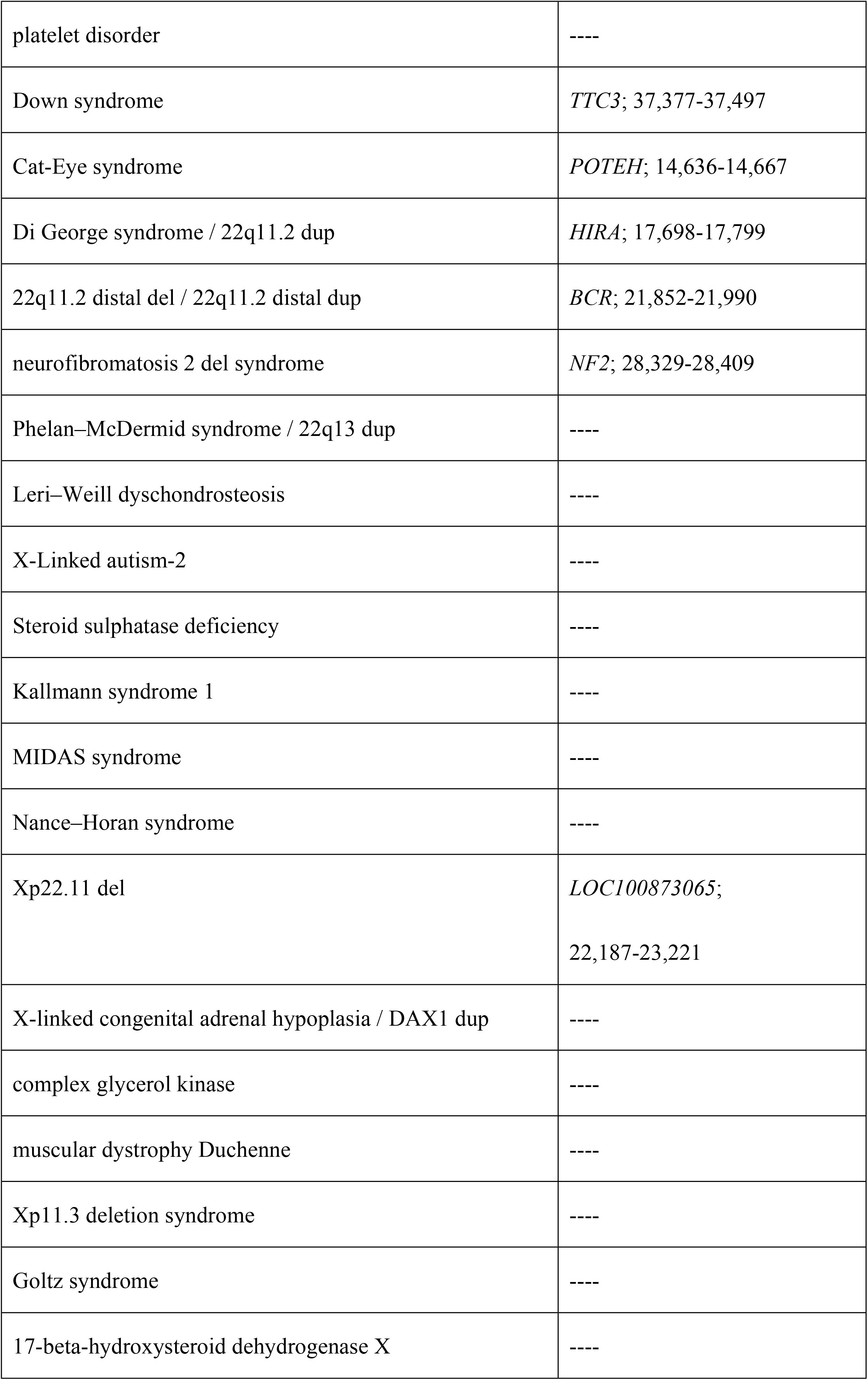
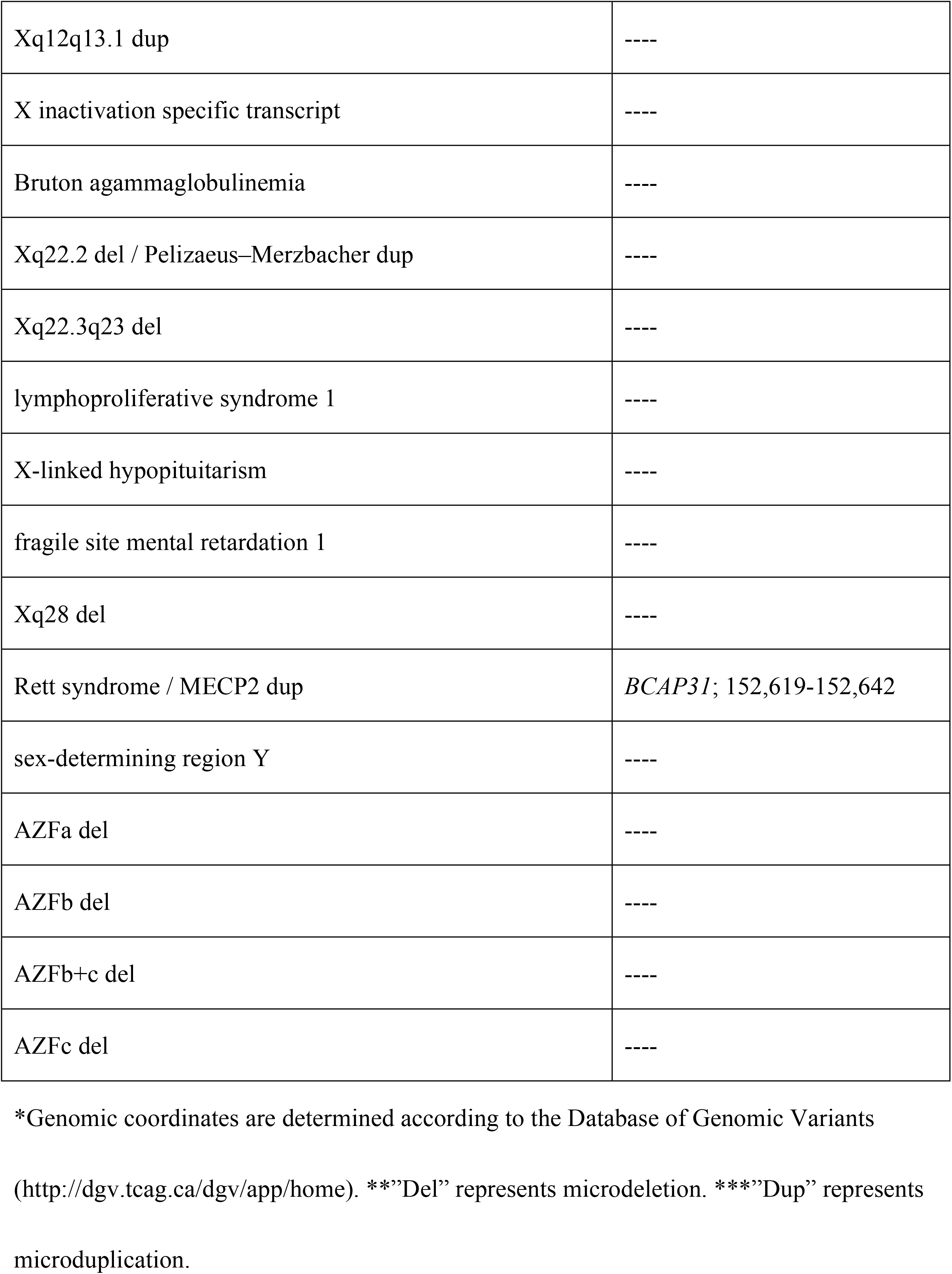
Microdeletion and microduplication syndromes and the featured ChIPed genes.

We only captured the genes related to 66.4%, rather than all, of the microdeletion and microduplication syndromes. Genes related to the remaining syndromes were not captured. Among many possible reasons that may be attributed to, one is that many deletion- and duplication-susceptible DNA segments might be epigenetically silenced due to the Jurkat cell-specific chromatin modification. These DNA segments are not accessible to TOP2. However, investigating the true reason is beyond the purpose and scope of the current study. Much more work is required to find the true attributing reasons which could be related to the ChIP-seq method, data analysis, statistics, cell biology, etc.

We did not either capture all the genes within the depleted and duplicated genomic DNA segment of any single syndrome. We only ChIPed those cells that were treated with etoposide for 5.5 hours. However, the repair of the TOP2-induced DNA break at any specific repair step often only involves a limited section of the DNA segment, we believe that a thorough kinetic and dynamic investigation of the repair process would identify more genetic sequences within the deleted or duplicated genomic segment.

### The poisoned TOP2-induced, γH2A.X-associated DNA was closely linked to many genetic sequences related to autistic spectrum disorders

To weigh how much the poisoned TOP2 would contribute to any specific types of the microdeletion and microduplication syndromes, we decided to investigate autism or autistic spectrum disorders (ASD) for the following reasons. Genes associated with ASD are often long, implicating that more topological strain is generated during transcription [38]. Moreover, the ASD-related genes are more thoroughly studied. We looked at three online ASD databases, Sfari Gene (https://gene.sfari.org/database/human-gene/), Autism Genetic Database (http://autism-genetic-db.net/browse/chromosome/y/feature/genes) and AutismK (http://autismkb.cbi.pku.edu.cn/ranked_gene.php). Of the 1051, 223 and 171 genes respectively reported by the databases, our ChIP captured 353 (33.6%), 70 (31.4%) and 46 (26.9%) of them (Supporting Information S1 Table). Therefore, the poisoned TOP2-induced, γH2A.X-associated DNA was closely linked to ASD.

### The poisoned TOP2-induced, γH2A.X-associated DNA was closely linked to many genetic sequences related to leukemia and other types of cancers

As discussed above, additional to the TOP2-poisoning anticancer drugs, the TOP2-poisoning bioflavonoids and pesticides have been linked to the chromosome translocation-related leukemia [7,13,16–18]. We then looked at whether the genes related to leukemia or other types of cancers were immunoprecipitated by our ChIP. Of the 138 genes that cause 132 types of fusions that were reviewed by Zhang and Rowley [39], we captured 55 (39.9%) genes that comprised 101 (76.5%) types of the fusions (Table 2.1 and Supporting Information S2 Table). We next looked at whether the poisoned TOP2 might also contribute to other types of cancers. Of the 82 oncogenes and 63 tumor suppressor genes that were defined according to the stringent criteria [40], our ChIP respectively pulled down 25 (30.5%) and 19 (30.2%) of them (Table 2.2 and Table 2.3). Therefore, the poisoned TOP2-induced, γH2A.X-associated DNA was closely linked to not only leukemia but also many types of cancers.

**Table 2.**
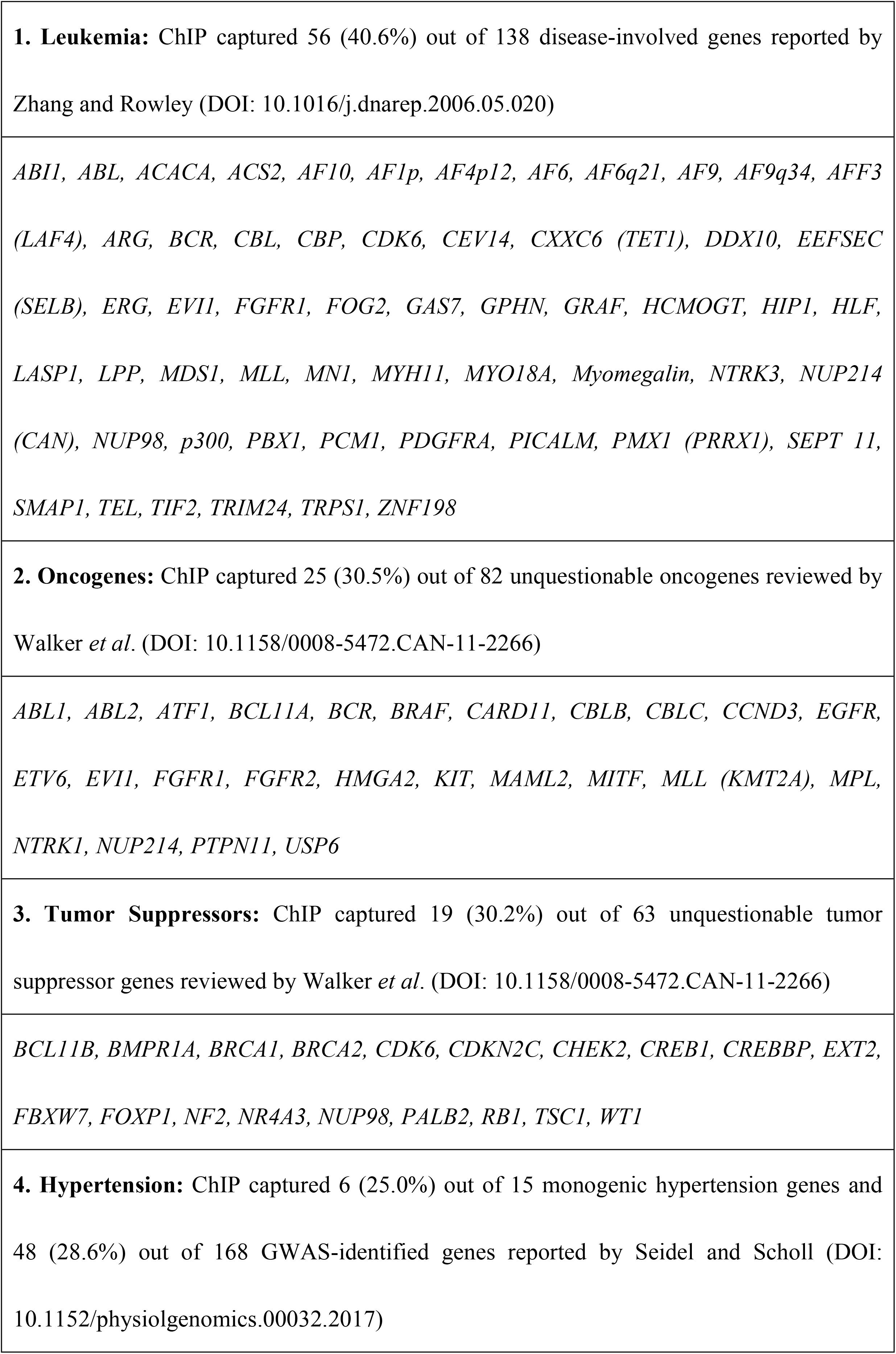
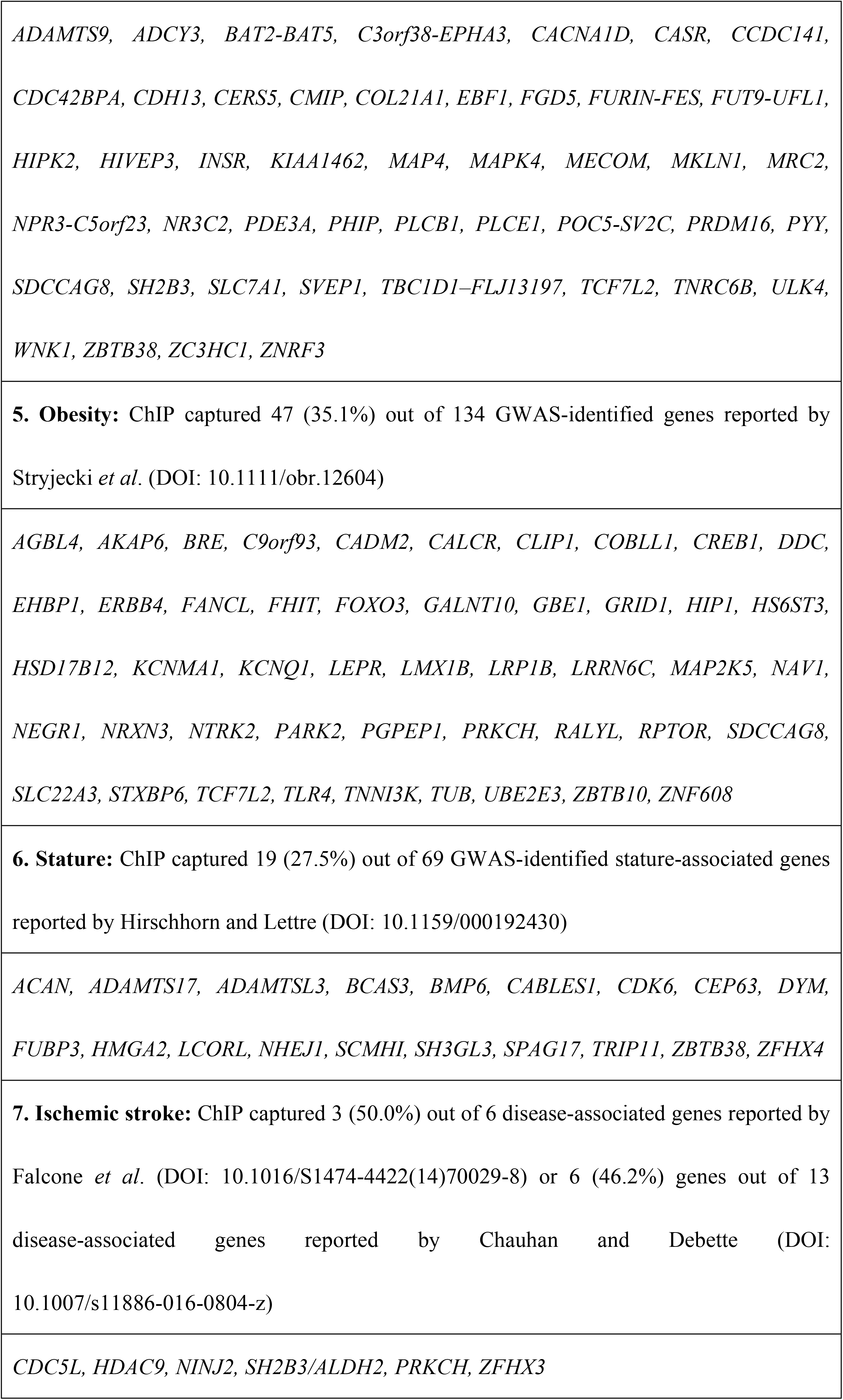
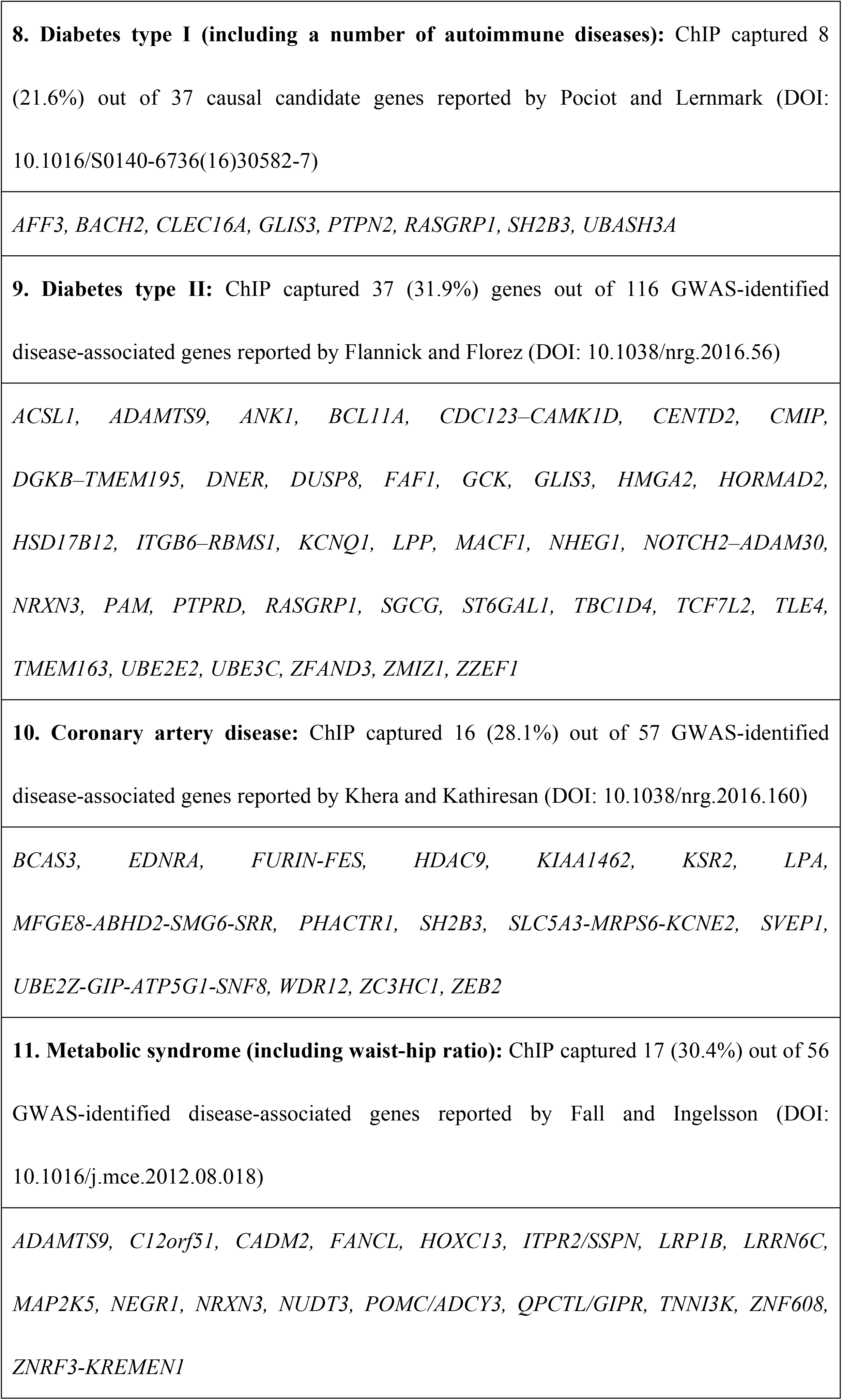
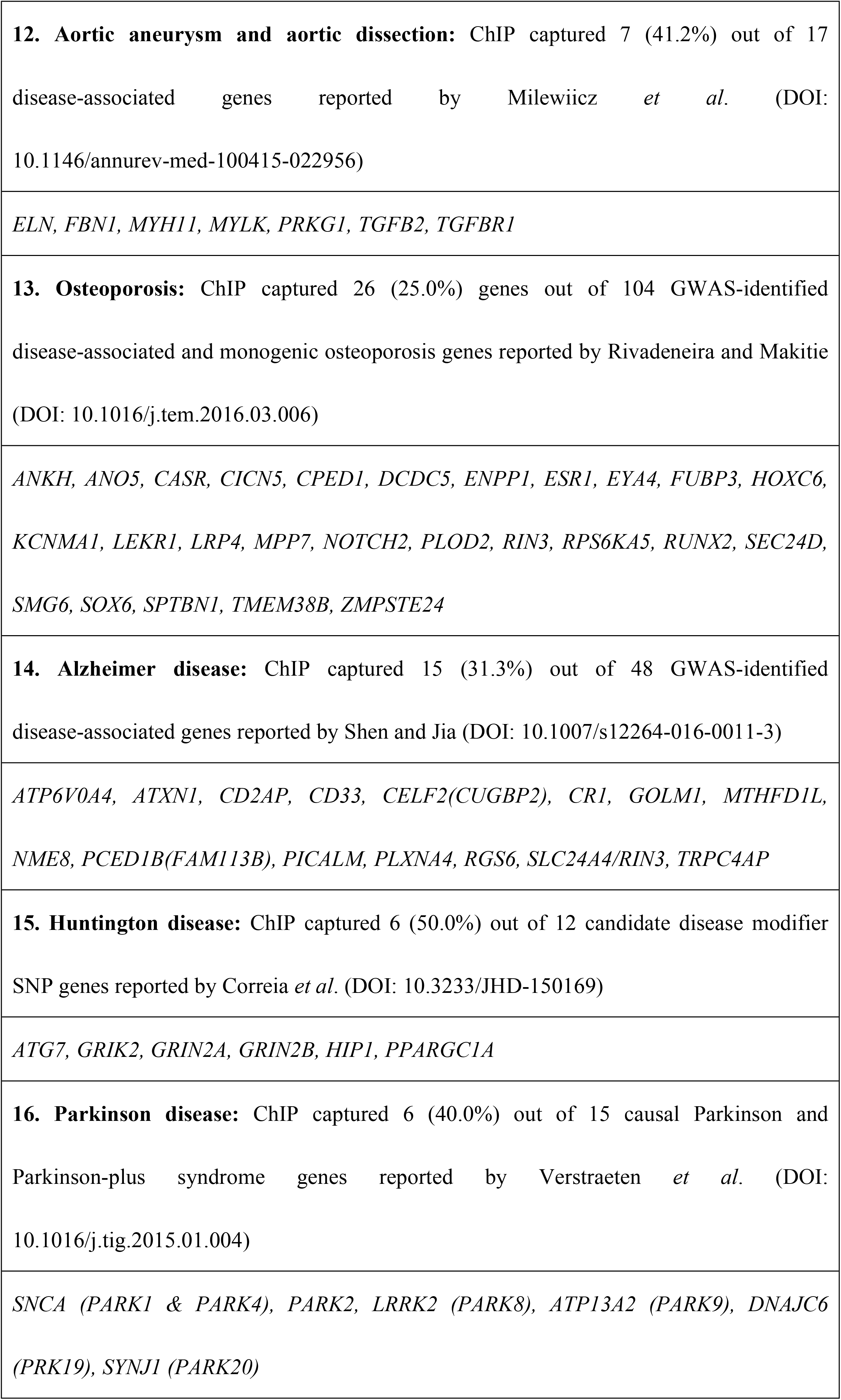
The disease-associated genes captured by the ChIP.

### The poisoned TOP2-induced, γH2A.X-associated DNA was closely linked to a large number of genetic sequences related to many common diseases or conditions

For the past several decades, the morbidity of many diseases and conditions, such as the cardiovascular and metabolic disorders, has been increased progressively and the onset age has been decreased continuously. On the other hand, the TOP2-poisoning bioflavonoids have been found to be associated with the genetic variations at the early or progressively later stage of a number of these diseases (such as atherosclerosis, cancer, and premature aging) [41]. To investigate whether the poisoned TOP2 might be linked to the genetic alterations of these diseases, we looked at the genes that have been reported to be associated with the other 13 common diseases and conditions, including hypertension, obesity, stature or height, ischemic stroke, type I diabetes, type II diabetes, coronary artery disease, metabolic syndrome (including hip-waist ratio), aortic aneurysm and dissection, osteoporosis, Alzheimer disease, Huntington disease, and Parkinson disease. Table 2.4 to Table 2.16 show that a large percent, ranging from about 25% to more than 50%, of the genes reported to be associated with these diseases or conditions were captured by our ChIP. Interestingly, except the *LPA* gene, all other coronary artery disease-associated genes captured by the ChIP were not related to lipid metabolism [42]. Likewise, most ChIP-captured stroke-associated genes were related to ischemic rather than hemorrhagic stroke [43]. One of the possibilities is that, for the Jurkat cells, the lipid metabolism- and hemorrhagic stroke-associated genes were epigenetically silenced or not located at the slow DNA replication region.

Additionally, we captured many genes, including *AFF3, BACH2, CLEC16A, GLIS3, PTPN2, RASGRP1, SH2B3, UBASH3A*, which contribute to both type I diabetes and other types of autoimmune diseases, supporting the early conclusion that these diseases share the same genetic etiology [44,45].

Overall, our data show that the poisoned TOP2-induced, γH2A.X-associated DNA was closely linked to many diseases.

## Discussion

TOP2 poisons are well known to cause various types of chromosome aberrations [6–8,14,15]. However, except a limited number of genes such as the *MLL* gene, most involved genes are not characterized, neither are their relationships with any specific diseases. Therefore, the one and only purpose of the current study is to identify the genetic sequences associated with the poisoned TOP2 and, particularly, its repair. To achieve our goal, we genome-widely immunoprecipitated the genomic segments that were associated with the γH2A.X induced by the TOP2 poison etoposide. As the signal of any γH2A.X-associated gene among a heterogeneous cell population is likely to be low, to capture as much amount of the signal as possible so that it fell into the sensitivity range of the ChIP-seq method, we treated the Jurkat cells with 100μM etoposide, waited for 5.5 hours when the γH2A.X was approximately at its peak level (Supporting Information S1 Fig) and then harvested the genomic DNA.

Our repeated ChIP experiments reproducibly captured 3707 transcriptable protein- and nonprotein-coding sequences. These sequences are likely to be TOP2-specific because they include 75.0% of the known topoisomerase-regulated genes essential for neurodevelopment (Supporting Information S3 Table) [46].

The TOP2-poisoning anticancer drugs, bioflavonoids and pesticides have been linked to leukemia and autism [7,13,16–18]. As expected, we did capture many genetic sequences related to leukemia and ASD (Table 2). However, unexpectedly, we pulled down many genetic sequences that belong with 66.4% of the CNV-related microdeletion and microduplication syndromes (Table 1). Moreover, we also identified a large number, ranging from 25% to 50%, of the genetic sequences whose CNVs and/or SNPs were associated with the other 16 common diseases or conditions that we investigated (Table 2). (Oncogenes and tumor suppressor genes belong to one disease, cancer.) These numbers are significant, considering that the human genome contains 154,484 to 323,827 transcriptable protein- and nonprotein-coding sequences [47] whereas we only ChIPed 3707 (1.1%-2.4%) of these sequences (NCBI SRA accession number SRP150381). Therefore, additional to the early reports that have linked TOP2 poisons to leukemia, autism, cardiovascular, and neurological development defects [48–50], our results raise the possibility that the poisoned TOP2 might also be linked to the genetic development of the aforementioned 17 diseases and conditions that we investigated. However, whether this linkage is of cause-effect type is yet to be confirmed.

Like most ChIP-seq studies, we only identified the genes that were related to the repair of the poisoned TOP2-induced DSB. Our study by no means directly addressed how these genetic sequences might mechanistically cause the CNVs and SNPs; investigating the biochemical or biological mechanism is not the purpose of the current study. However, at least for the *MLL* fusion, several mechanisms have been proposed [13,19–22,51]. While the poisoned TOP2 is considered to directly cause the *MLL* fusion [13], a number of studies have clearly demonstrated that the apoptotic CAD-induced high molecular weight (HMW) DNA cleavage contributes to its formation [19,20]. Indeed, γH2A.X as well as the non-homologous end joining DNA repair is activated by both poisoned TOP2- and CAD-induced DSBs [3,9,10,52,53]. Therefore, under our experiment conditions (i.e. 5.5 hours of the 100μM-etoposide treatment), a portion of the Jurkat cells might have entered apoptosis. As a result, our ChIPed genetic sequences could include those not only due to the initial poisoned TOP2-indcued DSBs prior to apoptosis, but also the DSBs resulting from the apoptotic HMW DNA cleavage. However, several pieces of evidence support our conclusion that the ChIPed genetic sequences are poisoned TOP2-specific. The evidence is as follows.

1. The nucleotide-resolution mapping of the TOP2- and CAD-mediated DSBs has found that neither TOP2 cleavage site nor apoptotic DNA cleavage site coincides with the *MLL* translocation hotspot [54].
2. TOP2 cleaves DNA at the anchorage site of the DNA loop domain, generating the apoptotic HMW DNA fragment, independent of the caspase-CAD pathway [55,56].
3. Binding of TOP2α to CAD greatly enhances TOP2α’s decatenation activity [57].
4. Simultaneous inhibiting caspase and poisoning TOP2α abolish the apoptotic chromosome condensation [57].

These early studies suggest that during apoptosis, TOP2 might work independently or cooperatively with CAD to cleave the looped DNA at the anchorage site, such as the TOP2 cleavage sequence-containing matrix attachment region [20,56,58.59], leading to the apoptotic HMW DNA fragmentation. As suggested by Betti *et al.*, TOP2 poisons might disrupt such roles of TOP2, causing the *MLL* fusion [60]. As a result, regardless whether the poisoned TOP2-induced *MLL* fusion is generated as a result of the initial DNA repair or subsequent apoptosis, it seems that the generation of the *MLL* fusion is related to the poisoned TOP2. Indeed, it has long been known 1) that many DNA damaging agents can induce CAD to cleave DNA within the breakpoint cluster region of the *MLL* gene [61] but, clinically, the therapy-related, *MLL* fusion-associated leukemia is almost always linked to the TOP2-poisoning anticancer drugs [23,62] and 2) that the formation of the *MLL* fusion is independent of the dosage of etoposide, as a wide range of doses, from 0.14-0.5μM, of etoposide can induce the fusion [36,37].

TOP2 is an enzyme that resolves the topological strain generated during DNA replication, transcription and other DNA transactions, meaning that TOP2 functions throughout the genome as long as the topological tension interferes with the DNA transactions. Therefore, concerns may be raised regarding 1) whether our ChIPed genetic sequences merely reflect the broad distribution of TOP2 in the genome and 2) whether there is any specificity in terms of the genomic location or disease linkage for the ChIPed genetic sequences over other general genetic sequences. Numerous studies and our findings support that the genomic location of the ChIPed genetic sequences, as well as their disease linkage, is not random but specific, which are described as follows.

1. The Human Genome Project estimates that the human genome has approximately 154,484 to 323,827 transcriptable protein- and nonprotein-coding sequences [47]. The number of the transcriptable sequences reproducibly pulled down by our ChIP are much fewer, only 3707 or between 1.1% to 2.4% of the total human sequences.
2. Of the diseases we investigated, we ChIPed 25% to more than 66% of the trranscriptable sequences that are associated with these diseases (Tables 1 and 2 and Supporting Information S1 and S2 Tables). If the sequences we ChIPed were not specific, average-wise, the number of the ChIPed disease-associated sequences would statically be close to 1.1%-2.4%, the percentage of the ChIPed (3707) over the total transcriptable sequences (154,484 to 323,827) in the genome.
3. The intensity of the topological strain, as well as the distribution of TOP2, across the genome or a transcribed and replicated gene is neither identical nor evenly distributed. Usually, more topological strain is accumulated during the transcription of the long genes such as those related to autism [38] and neurodevelopment [46]. Consistently, we ChIPed 33.6% of the autism-related genes (Supporting Information S1 Table) and 75.0% of the known topoisomerase-targeted neurodevelopment genes (Supporting Information S3 Table); both groups of genes are long [38,46].
4. The human genome contains various types of DNA repetitive and other noncanonical sequences [24,25] that may form non-B DNA secondary structures [63]. These noncanonical DNA sequences or non-B DNA structures are often found at the anchorage site of the looped DNA, such as the matrix or scaffold attachment site, or at the site where DNA transcription or replication is initiated or terminated. At these unique sites, DNA topoisomerases are required to unwind the DNA duplex or resolve the intensified DNA topological problem [6–8,63–70]. Consistently, Yu *et al.* have shown that the TOP2A cleavage sites are not evenly distributed throughout the genome [31]. Moreover, Bariar *et al.* have shown that it is the DNA sequences that drive the illegitimate DNA aberrations [71].
5. Most importantly, numerous studies have confirmed that the chromosome aberration breakpoints associated with CNVs often occur at these non-canonical DNA sequences and non-B DNA structures [63–70,72–74], where TOP2 is concentrated as described above.
6. Our ChIP pulled down 122 pseudogenes, 265 antisense genes and many genes that contain the major common fragile site (such as *ARHGAP15, CCSER1, CNTNAP2, CTNNA1, CTNNA3, DAB1, DLG2, DMD, DPYD, ESRRG, FHIT, GRID2, IL1RAPL1, IMMP2L, LARGE, LRP1B, MAD1L1, NBEA, PARK2, PDGFRA, PTPRC, RORA, SDK1, THSD7A, USH2A*, and *WWOX*) (NCBI SRA accession number SRP150381) [75] and early replicating fragile site (such as *BACH2*, *FOXP1*, *PVT1*, and *PRKCB*) [76]. The chromosome aberration-related DNA segmental duplication and DNA breakage often occur at these unique genetic sequences [29,75,76].

Accordingly, it is not surprising that our ChIPed genetic sequences specifically coincide with those associated with the CNV-related diseases such as the microdeletion and microduplication syndromes.

Additional to the genetic sequences related to the CNVs, we also captured many genetic sequences susceptible to the single nucleotide polymorphisms (SNPs), which is also consistent with the early findings that many disease-associated SNPs are often closely associated with the CNVs [77].

As a result, our data show that the repair, specifically γH2A.X activated by the poisoned TOP2 involves many genetic sequences that are linked to a variety of diseases. It should be noted that, we only investigated 17 diseases or conditions. However, our ChIPed genetic sequences involved much more diseases (NCBI SRA accession number SRP150381). For example, we ChIPed the *CFTR* gene whose mutation causes cystic fibrosis. Much more time is required to characterize the gene-disease relationships for all of the ChIPed genetic sequences.

We would like to emphasize that the only direct conclusion obtained from our data is that the repair of poisoned TOP2 is linked to the genetic sequences that are associated with many diseases. However, whether the linkage is of the etiology-disease (or cause-effect) type remains to be investigated. Currently, only a limited number of genes are known to directly contribute to certain diseases, such as the leukemia-associated *MLL* gene [62]. Nevertheless, the principle of our conclusion (i.e. the repair of poisoned TOP2 is linked to many disease-associated genes) should not be limited only to etoposide. It should also apply to other TOP2 poisons because what we have addressed is the repair of poisoned TOP2. The well-characterized etoposide is only used as a tool to poison TOP2.

Besides our direct conclusion, we also propose an hypothetical model which is describe in the following paragraphs. It should be noted that unlike the direct conclusion of our study, the model is extrapolated or deduced from not only our but also others’ data. Like most hypothetical models, it requires more work to be improved.

To interpret the clinical features of the CNV- and SNP-associated diseases, we propose a model according to our and others’ data which are as follows.

1. The TOP2-poisoning chemicals, such as the TOP2-poisoning bioflavonoids and pesticides, are widely distributed in the environment. Actually, a number of TOP2-poisoning bioflavonoids are recommended for the pregnant women as food supplements [50] or are contained in the infant formula [78].
2. A large number of previous studies have discovered that the environmental TOP2-poisoning chemicals, including the TOP2-poisoning bioflavonoids and pesticides, may contribute to a variety of human diseases [13–18,41,46,48–50,71,79–87]. More importantly, maternal ingestion of the TOP2-poisoning bioflavonoids has been linked to several developmental diseases that occur to the fetus and offspring [7,16–18,48–50].
3. Many laboratory studies have found that human and/or mouse monoploid spermocytes, oocytes and diploid embryonic cells are susceptible or related to the TOP2 poison-induced chromosome aberration [17,36,37,71,83–86,88].
4. Non-allelic homologous recombination, the most common DNA repair mechanism contributing to the CNV-related microdeletion and microduplication syndromes [29], is the major DNA repair pathway in meiosis during the maturation of monoploid spermocytes and oocytes. These monoploid germ cells have to use the non-allelic homologous recombination to repair the poisoned TOP2-induced DSB because, unlike the diploid cells, the monoploid cells do not have the allelic genes to conduct the copy-and-paste repair. Therefore, these cells are more prone to chromosome aberration [29].
5. Many studies have shown that the maternal exposure to the dietary bioflavonoids has a long-term effect on the offspring later in the adulthood [78].
6. The poisoned TOP2, including that caused by bioflavonoids, is known to cause chromosome translocation and other types of chromosome aberrations [6–8,13–15] although the involved genes have not been characterized; the latter is the purpose of the current study.
7. Ames have suggested that the dietary substances are more important than the environmental and occupational exposures to cause mutation [80,82].
8. Our data show that γH2A.X induced by the poisoned TOP2 is linked to many genetic sequences whose CNVs and SNPs are related to a variety of diseases (Tables 1 and 2 and Supporting Information S1 and S2 Tables).
9. Importantly, TOP2 is often located at the noncanonical DNA sequences or non-B DNA secondary structures which coincide with the breakage sites of various types of chromosome aberrations and the disease-associated CNVs [6–8,63–70].
10. Any cells, regardless of whether they are cells undergoing meiosis, embryonic cells, dividing cells or differentiated cells, require TOP2 to resolve the topological problem generated during the DNA transactions [6–8]. Therefore, they are prone to the mutation induced by TOP2 poisons.
11. The previous studies showing the co-occurrence of cancer and many CNV- or chromosome aberration-associated syndromes in children, back our model [89].

To summarize, the poisoned TOP2, on one hand, can cause chromosome aberration and, on the other hand, has been linked to a number of diseases. Moreover, the genomic site (repetitive and other non-canonical sequences) where TOP2 functions, coincides with the site where the chromosome aberration- and CNV-related breakage occurs. Therefore, the disease-associated CNVs and SNPs might be specifically related to the poisoned TOP2, just like the *MLL* fusion that contributes to the pediatric and therapy-related leukemia. Monoploid germ cells are particularly vulnerable because they do not have the allelic genes to conduct the copy-and-paste repair.

Fig 1 (Please refer to the end of the manuscript for the caption of Fig 1.) shows our model which seems able to explain many clinical features, including mosaicism, two-hit theory, polygenic traits, pleiotropy, and organ- and age-specificity, of both microdeletion and microduplication syndromes and CNV-associated diseases.

**Fig 1.**
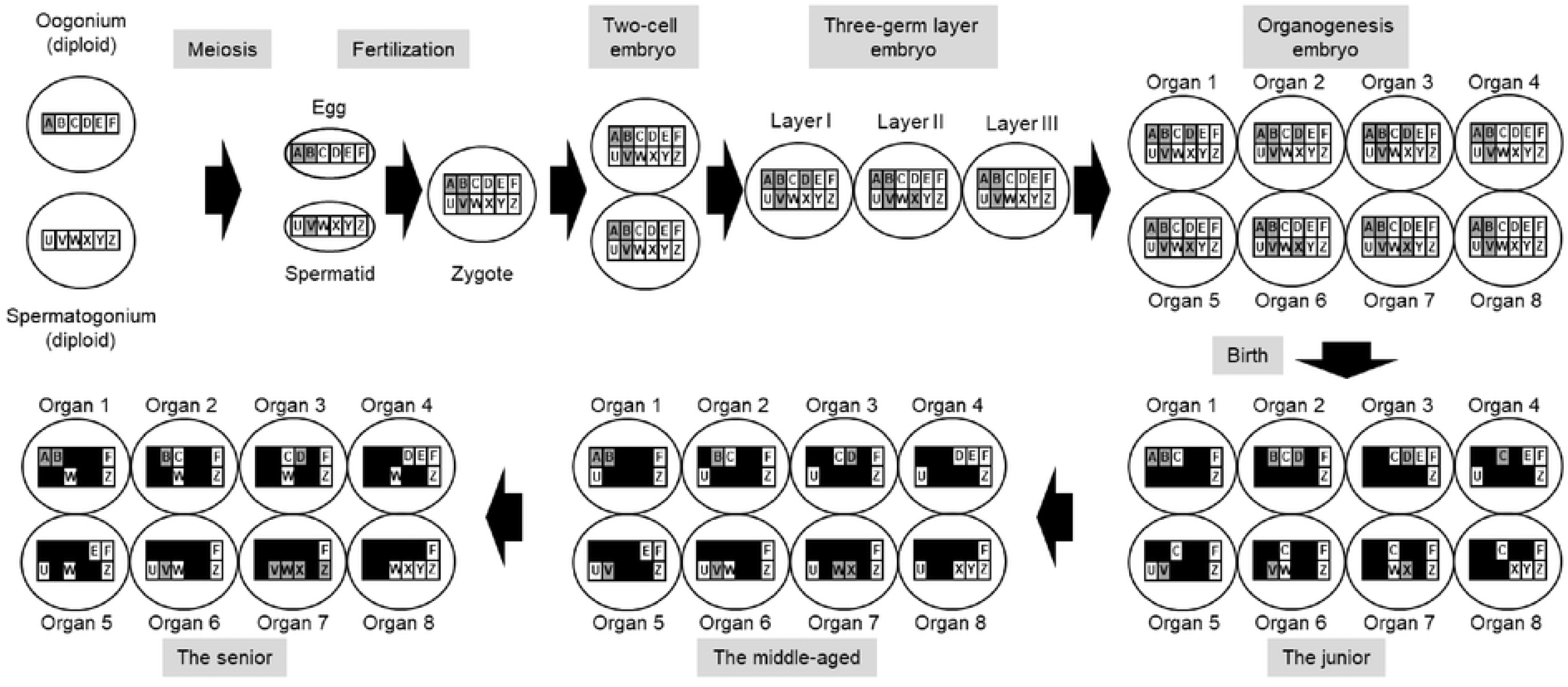
A hypothetical model of the CNV- and SNP-related diseases. Schematically presented is the maturation process of the parental germ cells and the development of a human being from a fertilized egg (including the two-cell embryo, three germ-cell layer embryo and embryo undergoing organogenesis) to a senior adult. At any developmental stages, the cells are prone to the mutation caused by a TOP2 poison. The numbered large ovals represent various human organs such as the lungs. The lettered small square boxes represent the genes. Letters A through Z within the small square boxes represent the gene names. Only one set of the alleles are illustrated. An open square box shows a wild-type gene that is expressed at that specific development stage. A hatched square box represents a mutated gene that is expressed at that specific development stage. Its first appearance indicates that it is initially mutated at that development stage. A closed square box indicates a silenced gene. Organ 1, 2 and 3 developed from layer I of the three-layer embryo. Organ 5, 6 and 7 developed from layer II of the three-layer embryo. Organ 4 and 8 developed from layer III of the three-layer embryo. The development stage- and organ-specific exposure to the TOP2 poison determines when and where a gene is mutated. Here, the TOP2 poison-induced genetic mutations occur as follows: Gene A before meiosis (inherited), Genes B and V during meiosis (*de novo*), Genes D (layer I of the three-layer embryo) and X (layer II of the three-layer embryo) at the three-germ layer embryo stage (*de novo*), Genes C, W and Z of Organs 4, 7 and 7 at the junior, middle-aged and senior stages, respectively. If a gene is mutated during meiosis or at the one-cell embryo stage, all body cells harbor the mutation (Genes B and V). If the gene of only a portion of the body is mutated after the one-cell embryo stage, mosaicism ensues (Genes D, X, C, W, and Z). A monogenic disease of Organ 3 develops if a mutated Gene D is sufficient to cause the disease. The polygenic, “two-hit” or modifier-requiring disease of Organ 7 requires all Genes V, W, X, and Z to be mutated accumulatively at several development stages. Pleiotropy occurs when the mutated Gene B causes both diseases of Organ 1 and Organ 2.

The morbidity of many aforementioned diseases, such as ASD and diabetes, have been increased progressively, particularly in the industrialized regions. However, the etiologies for the genetic development of the diseases are elusive. On the other hand, the TOP2-poisoning bioflavonoids are almost ingested daily as food components, additives (in the processed food), supplements, or contaminators (pesticides), often at the high doses that have long been known to cause mutation and genome instability [79,80]. For example, the daily dose of the bioflavonoids recommended by the manufactures can be as high as 1 to 2 grams per day. At these doses, the plasma levels can be 10 to 20 times higher than those that can induce mouse *MLL* fusion [81]. Besides, bioflavonoids at these doses have been linked to a number of human diseases [7,13,16–18,48–50]. Therefore, a large number of studies have proposed that the dietary TOP2-poisoning bioflavonoids clinically contribute to the development of many diseases, including leukemia, autism, developmental diseases, and neurological defects [7,13,18,41,46,48–50,71,78–87]. Our data that the repair of the poisoned TOP2 involves a large number of disease-associated genes, support this theory. Moreover, according to our and, particularly, others’ data as described above, we extend the theory to include many other diseases and propose that TOP2 poisons, including the TOP2-poisoning bioflavonoids, might also be an important, clinically relevant DNA-mutating etiology for other CNV- and SNP-related diseases. If the hypothesis is confirmed, the morbidity rates of many diseases can be decreased by reducing human’s exposure to the TOP2-poisoing substances.

It should be noted that only the B- and C-ring dietary bioflavonoids have TOP2 poisoning activities [87]. Moreover, various TOP2-poisiong bioflavonoids exhibit disparate potency and efficacy [71]. Additionally, multiple cellular effects of bioflavonoids have been observed, including antioxidant, pro-oxidant, anti-mutagenic, mutagenic, carcinogenic, and anti-carcinogenic activities among many others [79]. Being antioxidant, antimutagenic and anticarcinogenic, bioflavonoids also benefit the health for adults [79]. Therefore, more studies, including modifying the B- and C-ring of the dietary bioflavonoids, might be warranted for their effects on human beings, particularly pregnant women and infants.

Lastly, we hope that any researchers who are interested and are also funded can further the investigation. Afterall, the development disorders, particularly autism, affect many children across the world, particularly in the industrialized regions where the processed food is routinely consumed.

In conclusion, we have found that many genetic sequences associated with the repair of the poisoned TOP2, are the candidates for not only leukemia and ASD but also a variety of other CNV- and SNP-related diseases. Our results raise the possibility that the TOP2-poisoining chemicals such as the dietary bioflavonoids might be involved in the genetic development of many diseases. Additionally, according to our and, particularly, others’ data, we propose a yet-to-be-confirmed hypothetical model that seems able to interpret many features of the CNV-related microdeletion and microduplication syndromes (Figure 1). Future cell studies using the TOP2-poisoning bioflavonoids, animal experiments and human epidemiolocal studies are required to validate the model.

## Materials and methods

### Cell culture and drug treatment

Human T-cell acute lymphoblastic leukemia Jurkat cells were kindly provided by the Stem Cell Bank of Chinese Academy of Sciences. These Jurkat cells were routinely maintained in the RPMI 1640 media (Gibco). They were mock-treated with 0.1% DMSO (Cat# D2438, Sigma-Aldrich) or treated with 100μM etoposide (Cat# E1383, Sigma-Aldrich) that was prepared from the 100mM stock solution in DMSO.

### Western blot

Cells were collected after a series of time intervals following the continuous treatment of etoposide. Western blot analysis was performed as descried previously [30] except that the antibodies against γH2A.X (Cat# 07-164, Millipore) were used as the primary antibodies at the 1:2000 dilution of manufacture’s stock solution.

### ChIP-seq

ChIP was performed using the EZ-ChIP™ Chromatin Immunoprecipitation Kit (Cat# 17-371, Millipore) according to manufacturer’s instructions. Both DMSO mock-treated control and etoposide-treated cells were crosslinked with formaldehyde (Cat# F8775, Sigma-Aldrich) at room temperature for 10 minutes. Sonication of the formaldehyde-crosslinked cells was conducted using the Covaris S220 sonicator (Gene Company Limited) at the peak power of 140.0, duty factor of 5.0 and cycle/burst of 200 for 600 seconds. The antibodies against γH2A.X (Cat# 07-164, Millipore), at the 1:250 dilution from manufacture’s stock, were used to immunoprecipitate the etoposide-induced DNA repair complexes. To increase the specificity and reproducibility of the peaks identified by DNA sequencing, 20 to 30 nanograms of DNA from the samples of not only a single ChIP experiment but also a pool of 8 ChIP experiments were subjected to the DNA sequencing. The sequencing libraries were prepared following manufacturer’s instructions of the SPARK DNA Sample Prep Kit (Cat# SPK0001-V08, Enzymatics). The purified DNA was sequenced paired-end with a read-length of 150 bases on an Illumina HiSeq X Ten sequencer at Shanghai Biotechnology Corporation. The ChIP-seq reads were mapped to the human genome (hg18) with Bowtie2 using the default parameters. The statistically significant peaks (*P*<0.01) were called using MACS (version 1.4.2) with the default parameters and according to the standard protocols. Both DNA of the DMSO mock-treated cells and input DNA were used for the peak identification of the etoposide-treated samples. Only the peaks that were reproducibly identified from the repeated experiments were annotated to the nearest genes using the NCBI and UCSC databases. The sequencing data were deposited at the NCBI SRA under the accession number SRP150381.

## Acknowledgements

Not applicable.

## Supporting Information Captions

**S1 Fig. The kinetics of the etoposide-induced γH2A.X by the western blot analysis**

**S1 Table. The ChIP-captured autism-related genes reported by 3 databases**

**S2 Table. The leukemia chromosome rearrangement-involved genes captured by ChIP**

**S3 Table. The topoisomerase-regulated neurodevelopment-related genes captured by ChIP**

## References

1. Rogakou EP, Pilch DR, Orr AH, Ivanova VS, Bonner WM. DNA double-stranded breaks induce histone H2AX phosphorylation on serine 139. J Biol Chem 1998 Mar 6;273(10):5858–5868.

2. Kinner A, Wu W, Staudt C, Iliakis G. Gamma-H2AX in recognition and signaling of DNA double-strand breaks in the context of chromatin. Nucleic Acids Res 2008 Oct;36(17):5678–5694.

3. Mishima M. Chromosomal aberrations, clastogens vs aneugens. Front Biosci (Schol Ed) 2017 Jan 1;9:1–16.

4. Georgoulis A, Vorgias CE, Chrousos GP, Rogakou EP. Genome Instability and gammaH2AX. Int J Mol Sci 2017 Sep 15;18(9):10.3390/ijms18091979.

5. Seo J, Kim SC, Lee HS, Kim JK, Shon HJ, Salleh NL, et al. Genome-wide profiles of H2AX and gamma-H2AX differentiate endogenous and exogenous DNA damage hotspots in human cells. Nucleic Acids Res 2012 Jul;40(13):5965–5974.

6. Borde V, Duguet M. The mapping of DNA topoisomerase sites in vivo: a tool to enlight the functions of topoisomerases. Biochimie 1998 Mar;80(3):223–233.

7. Deweese JE, Osheroff N. The DNA cleavage reaction of topoisomerase II: wolf in sheep’s clothing. Nucleic Acids Res 2009 Feb;37(3):738–748.

8. Pommier Y, Sun Y, Huang SN, Nitiss JL. Roles of eukaryotic topoisomerases in transcription, replication and genomic stability. Nat Rev Mol Cell Biol 2016 Nov;17(11):703–721.

9. Darzynkiewicz Z, Halicka DH, Tanaka T. Cytometric assessment of DNA damage induced by DNA topoisomerase inhibitors. Methods Mol Biol 2009;582:145–153.

10. de Campos-Nebel M, Larripa I, Gonzalez-Cid M. Topoisomerase II-mediated DNA damage is differently repaired during the cell cycle by non-homologous end joining and homologous recombination. PLoS One 2010 Sep 2;5(9):10.1371/journal.pone.0012541.

11. Bromberg KD, Burgin AB, Osheroff N. A two-drug model for etoposide action against human topoisomerase IIalpha. J Biol Chem 2003 Feb 28;278(9):7406–7412.

12. Muslimovic A, Nystrom S, Gao Y, Hammarsten O. Numerical analysis of etoposide induced DNA breaks. PLoS One 2009 Jun 10;4(6):e5859.

13. Strick R, Strissel PL, Borgers S, Smith SL, Rowley JD. Dietary bioflavonoids induce cleavage in the MLL gene and may contribute to infant leukemia. Proc Natl Acad Sci U S A 2000 Apr 25;97(9):4790–4795.

14. Ferguson LR, Baguley BC. Topoisomerase II enzymes and mutagenicity. Environ Mol Mutagen 1994;24(4):245–261.

15. Baguley BC, Ferguson LR. Mutagenic properties of topoisomerase-targeted drugs. Biochim Biophys Acta 1998 Oct 1;1400(1-3):213–222.

16. Roman GC. Autism: transient in utero hypothyroxinemia related to maternal flavonoid ingestion during pregnancy and to other environmental antithyroid agents. J Neurol Sci 2007 Nov 15;262(1-2):15–26.

17. Hernandez AF, Menendez P. Linking Pesticide Exposure with Pediatric Leukemia: Potential Underlying Mechanisms. Int J Mol Sci 2016 Mar 29;17(4):461.

18. Bakian AV, VanDerslice JA. Pesticides and autism. BMJ 2019 Mar 20;364:l1149.

19. Stanulla M, Wang J, Chervinsky DS, Thandla S, Aplan PD. DNA cleavage within the MLL breakpoint cluster region is a specific event which occurs as part of higher-order chromatin fragmentation during the initial stages of apoptosis. Mol Cell Biol 1997 Jul;17(7):4070–4079.

20. Hars ES, Lyu YL, Lin CP, Liu LF. Role of apoptotic nuclease caspase-activated DNase in etoposide-induced treatment-related acute myelogenous leukemia. Cancer Res 2006 Sep 15;66(18):8975–8979.

21. Gole B, Wiesmuller L. Leukemogenic rearrangements at the mixed lineage leukemia gene (MLL)-multiple rather than a single mechanism. Front Cell Dev Biol 2015 Jun 25;3:41.

22. Gothe HJ, Bouwman BAM, Gusmao EG, Piccinno R, Petrosino G, Sayols S, et al. Spatial Chromosome Folding and Active Transcription Drive DNA Fragility and Formation of Oncogenic MLL Translocations. Mol Cell 2019 Jul 25;75(2):267–283.e12.

23. Wright RL, Vaughan AT. A systematic description of MLL fusion gene formation. Crit Rev Oncol Hematol 2014 Sep;91(3):283–291.

24. Girirajan S, Eichler EE. Phenotypic variability and genetic susceptibility to genomic disorders. Hum Mol Genet 2010 Oct 15;19(R2):R176–87.

25. Harel T, Lupski JR. Genomic disorders 20 years on-mechanisms for clinical manifestations. Clin Genet 2018 Mar;93(3):439–449.

26. Deak KL, Horn SR, Rehder CW. The evolving picture of microdeletion/microduplication syndromes in the age of microarray analysis: variable expressivity and genomic complexity. Clin Lab Med 2011 Dec;31(4):543–64, viii.

27. Weise A, Mrasek K, Klein E, Mulatinho M, Llerena JC,Jr, Hardekopf D, et al. Microdeletion and microduplication syndromes. J Histochem Cytochem 2012 May;60(5):346–358.

28. Watson CT, Marques-Bonet T, Sharp AJ, Mefford HC. The genetics of microdeletion and microduplication syndromes: an update. Annu Rev Genomics Hum Genet 2014;15:215–244.

29. Carvalho CM, Lupski JR. Mechanisms underlying structural variant formation in genomic disorders. Nat Rev Genet 2016 Apr;17(4):224–238.

30. Yang L, Dong F, Yang Q, Yang PF, Wu R, Wu QF, et al. FGF13 Selectively Regulates Heat Nociception by Interacting with Nav1.7. Neuron 2017 Feb 22;93(4):806–821.e9.

31. Yu X, Davenport JW, Urtishak KA, Carillo ML, Gosai SJ, Kolaris CP, et al. Genome-wide TOP2A DNA cleavage is biased toward translocated and highly transcribed loci. Genome Res 2017 Jul;27(7):1238–1249.

32. Dellino GI, Palluzzi F, Chiariello AM, Piccioni R, Bianco S, Furia L, et al. Release of paused RNA polymerase II at specific loci favors DNA double-strand-break formation and promotes cancer translocations. Nat Genet 2019 Jun;51(6):1011–1023.

33. Mao Y, Desai SD, Ting CY, Hwang J, Liu LF. 26 S proteasome-mediated degradation of topoisomerase II cleavable complexes. J Biol Chem 2001 Nov 2;276(44):40652–40658.

34. Alchanati I, Teicher C, Cohen G, Shemesh V, Barr HM, Nakache P, et al. The E3 ubiquitin-ligase Bmi1/Ring1A controls the proteasomal degradation of Top2alpha cleavage complex - a potentially new drug target. PLoS One 2009 Dec 1;4(12):e8104.

35. Ban Y, Ho CW, Lin RK, Lyu YL, Liu LF. Activation of a novel ubiquitin-independent proteasome pathway when RNA polymerase II encounters a protein roadblock. Mol Cell Biol 2013 Oct;33(20):4008–4016.

36. Bueno C, Catalina P, Melen GJ, Montes R, Sanchez L, Ligero G, et al. Etoposide induces MLL rearrangements and other chromosomal abnormalities in human embryonic stem cells. Carcinogenesis 2009 Sep;30(9):1628–1637.

37. Moneypenny CG, Shao J, Song Y, Gallagher EP. MLL rearrangements are induced by low doses of etoposide in human fetal hematopoietic stem cells. Carcinogenesis 2006 Apr;27(4):874–881.

38. King IF, Yandava CN, Mabb AM, Hsiao JS, Huang HS, Pearson BL, et al. Topoisomerases facilitate transcription of long genes linked to autism. Nature 2013 Sep 5;501(7465):58–62.

39. Zhang Y, Rowley JD. Chromatin structural elements and chromosomal translocations in leukemia. DNA Repair (Amst) 2006 Sep 8;5(9-10):1282–1297.

40. Walker EJ, Zhang C, Castelo-Branco P, Hawkins C, Wilson W, Zhukova N, et al. Monoallelic expression determines oncogenic progression and outcome in benign and malignant brain tumors. Cancer Res 2012 Feb 1;72(3):636–644.

41. Ferguson LR, Philpott M. Nutrition and mutagenesis. Annu Rev Nutr 2008;28:313–329.

42. Khera AV, Kathiresan S. Genetics of coronary artery disease: discovery, biology and clinical translation. Nat Rev Genet 2017 Jun;18(6):331–344.

43. Chauhan G, Debette S. Genetic Risk Factors for Ischemic and Hemorrhagic Stroke. Curr Cardiol Rep 2016 Dec;18(12):124-016-0804-z.

44. Roizen JD, Bradfield JP, Hakonarson H. Progress in understanding type 1 diabetes through its genetic overlap with other autoimmune diseases. Curr Diab Rep 2015 Nov;15(11):102-015-0668-4.

45. Pociot F, Lernmark A. Genetic risk factors for type 1 diabetes. Lancet 2016 Jun 4;387(10035):2331–2339.

46. Vokalova L, Durdiakova J, Ostatnikova D. Topoisomerases interlink genetic network underlying autism. Int J Dev Neurosci 2015 Dec;47(Pt B):361–368.

47. Salzberg SL. Open questions: How many genes do we have? BMC Biol 2018 Aug 20;16(1):94-018-0564-x.

48. Zielinsky P, Piccoli AL,Jr, Manica JL, Nicoloso LH, Menezes H, Busato A, et al. Maternal consumption of polyphenol-rich foods in late pregnancy and fetal ductus arteriosus flow dynamics. J Perinatol 2010 Jan;30(1):17–21.

49. Hahn M, Baierle M, Charao MF, Bubols GB, Gravina FS, Zielinsky P, et al. Polyphenol-rich food general and on pregnancy effects: a review. Drug Chem Toxicol 2017 Jul;40(3):368–374.

50. Barenys M, Masjosthusmann S, Fritsche E. Is Intake of Flavonoid-Based Food Supplements During Pregnancy Safe for the Developing Child? A Literature Review. Curr Drug Targets 2017;18(2):196–231.

51. Gomez-Herreros F. DNA Double Strand Breaks and Chromosomal Translocations Induced by DNA Topoisomerase II. Front Mol Biosci 2019 Dec 10;6:141.

52. Gomez-Herreros F, Romero-Granados R, Zeng Z, Alvarez-Quilon A, Quintero C, Ju L, et al. TDP2-dependent non-homologous end-joining protects against topoisomerase II-induced DNA breaks and genome instability in cells and in vivo. PLoS Genet 2013;9(3):e1003226.

53. Rogakou EP, Nieves-Neira W, Boon C, Pommier Y, Bonner WM. Initiation of DNA fragmentation during apoptosis induces phosphorylation of H2AX histone at serine 139. J Biol Chem 2000 Mar 31;275(13):9390–9395.

54. Mirault ME, Boucher P, Tremblay A. Nucleotide-resolution mapping of topoisomerase-mediated and apoptotic DNA strand scissions at or near an MLL translocation hotspot. Am J Hum Genet 2006 Nov;79(5):779–791.

55. Li TK, Chen AY, Yu C, Mao Y, Wang H, Liu LF. Activation of topoisomerase II-mediated excision of chromosomal DNA loops during oxidative stress. Genes Dev 1999 Jun 15;13(12):1553–1560.

56. Solovyan VT, Bezvenyuk ZA, Salminen A, Austin CA, Courtney MJ. The role of topoisomerase II in the excision of DNA loop domains during apoptosis. J Biol Chem 2002 Jun 14;277(24):21458–21467.

57. Durrieu F, Samejima K, Fortune JM, Kandels-Lewis S, Osheroff N, Earnshaw WC. DNA topoisomerase IIalpha interacts with CAD nuclease and is involved in chromatin condensation during apoptotic execution. Curr Biol 2000 Jul 27-Aug 10;10(15):923–926.

58. Khodarev NN, Bennett T, Shearing N, Sokolova I, Koudelik J, Walter S, et al. LINE L1 retrotransposable element is targeted during the initial stages of apoptotic DNA fragmentation. J Cell Biochem 2000 Sep 7;79(3):486–495.

59. Kantidze OL, Iarovaia OV, Razin SV. Assembly of nuclear matrix-bound protein complexes involved in non-homologous end joining is induced by inhibition of DNA topoisomerase II. J Cell Physiol 2006 Jun;207(3):660–667.

60. Betti CJ, Villalobos MJ, Diaz MO, Vaughan AT. Apoptotic stimuli initiate MLL-AF9 translocations that are transcribed in cells capable of division. Cancer Res 2003 Mar 15;63(6):1377–1381.

61. Sim SP, Liu LF. Nucleolytic cleavage of the mixed lineage leukemia breakpoint cluster region during apoptosis. J Biol Chem 2001 Aug 24;276(34):31590–31595.

62. Rashidi A, Fisher SI. Therapy-related acute promyelocytic leukemia: a systematic review. Med Oncol 2013;30(3):625-013-0625-5. Epub 2013 Jun 15.

63. Wells RD. Discovery of the role of non-B DNA structures in mutagenesis and human genomic disorders. J Biol Chem 2009 Apr 3;284(14):8997–9009.

64. Bose P, Hermetz KE, Conneely KN, Rudd MK. Tandem repeats and G-rich sequences are enriched at human CNV breakpoints. PLoS One 2014 Jul 1;9(7):e101607.

65. Kantidze OL, Razin SV. Chromatin loops, illegitimate recombination, and genome evolution. Bioessays 2009 Mar;31(3):278–286.

66. Usdin K, House NC, Freudenreich CH. Repeat instability during DNA repair: Insights from model systems. Crit Rev Biochem Mol Biol 2015 Mar-Apr;50(2):142–167.

67. Rothschild G, Basu U. Lingering Questions about Enhancer RNA and Enhancer Transcription-Coupled Genomic Instability. Trends Genet 2017 Feb;33(2):143–154.

68. Hall AC, Ostrowski LA, Pietrobon V, Mekhail K. Repetitive DNA loci and their modulation by the non-canonical nucleic acid structures R-loops and G-quadruplexes. Nucleus 2017 Mar 4;8(2):162–181.

69. Vasquez KM, Wang G. The yin and yang of repair mechanisms in DNA structure-induced genetic instability. Mutat Res 2013 Mar-Apr;743-744:118–131.

70. Canela A, Maman Y, Jung S, Wong N, Callen E, Day A, et al. Genome Organization Drives Chromosome Fragility. Cell 2017 Jul 27;170(3):507–521.e18.

71. Bariar B, Vestal CG, Deem B, Goodenow D, Ughetta M, Engledove RW, et al. Bioflavonoids promote stable translocations between MLL-AF9 breakpoint cluster regions independent of normal chromosomal context: Model system to screen environmental risks. Environ Mol Mutagen 2019 Mar;60(2):154–167.

72. Cooper GM, Nickerson DA, Eichler EE. Mutational and selective effects on copy-number variants in the human genome. Nat Genet 2007 Jul;39(7 Suppl):S22–9.

73. Conrad DF, Pinto D, Redon R, Feuk L, Gokcumen O, Zhang Y, et al. Origins and functional impact of copy number variation in the human genome. Nature 2010 Apr 1;464(7289):704–712.

74. van Binsbergen E. Origins and breakpoint analyses of copy number variations: up close and personal. Cytogenet Genome Res 2011;135(3-4):271–276.

75. Gao G, Smith DI. Very large common fragile site genes and their potential role in cancer development. Cell Mol Life Sci 2014 Dec;71(23):4601–4615.

76. Barlow JH, Faryabi RB, Callen E, Wong N, Malhowski A, Chen HT, et al. Identification of early replicating fragile sites that contribute to genome instability. Cell 2013 Jan 31;152(3):620–632.

77. McCarroll SA. Extending genome-wide association studies to copy-number variation. Hum Mol Genet 2008 Oct 15;17(R2):R135–42.

78. Vanhees K, Vonhogen IG, van Schooten FJ, Godschalk RW. You are what you eat, and so are your children: the impact of micronutrients on the epigenetic programming of offspring. Cell Mol Life Sci 2014 Jan;71(2):271–285.

79. Skibola CF, Smith MT. Potential health impacts of excessive flavonoid intake. Free Radic Biol Med 2000 Aug;29(3-4):375–383.

80. Ferguson LR. Role of plant polyphenols in genomic stability. Mutat Res 2001 Apr 18;475(1-2):89–111.

81. Vanhees K, Coort S, Ruijters EJ, Godschalk RW, van Schooten FJ, Barjesteh van Waalwijk van Doorn-Khosrovani S. Epigenetics: Prenatal exposure to genistein leaves a permanent signature on the hematopoietic lineage. FASEB J 2011 Feb;25(2):797–807

82. Ames BN. Identifying environmental chemicals causing mutations and cancer. Science 1979 May 11;204(4393):587–593.

83. Ross JA, Potter JD, Reaman GH, Pendergrass TW, Robison LL. Maternal exposure to potential inhibitors of DNA topoisomerase II and infant leukemia (United States): a report from the Children’s Cancer Group. Cancer Causes Control 1996 Nov;7(6):581–590.

84. Barjesteh van Waalwijk van Doorn-Khosrovani,S., Janssen J, Maas LM, Godschalk RW, Nijhuis JG, van Schooten FJ. Dietary flavonoids induce MLL translocations in primary human CD34+ cells. Carcinogenesis 2007 Aug;28(8):1703–1709.

85. Pelkonen O, Terron A, Hernandez AF, Menendez P, Bennekou SH, EFSA WG EPI1 and its other members. Chemical exposure and infant leukaemia: development of an adverse outcome pathway (AOP) for aetiology and risk assessment research. Arch Toxicol 2017 Aug;91(8):2763–2780.

86. Thys RG, Lehman CE, Pierce LC, Wang YH. Environmental and chemotherapeutic agents induce breakage at genes involved in leukemia-causing gene rearrangements in human hematopoietic stem/progenitor cells. Mutat Res 2015 Sep;779:86–95.

87. Bandele OJ, Clawson SJ, Osheroff N. Dietary polyphenols as topoisomerase II poisons: B ring and C ring substituents determine the mechanism of enzyme-mediated DNA cleavage enhancement. Chem Res Toxicol 2008 Jun;21(6):1253–1260.

88. Mailhes JB, Marchetti F, Young D, London SN. Numerical and structural chromosome aberrations induced by etoposide (VP16) during oocyte maturation of mice: transmission to one-cell zygotes and damage to dictyate oocytes. Mutagenesis 1996 Jul;11(4):357–361.

89. Zimmerman R, Schimmenti L, Spector L. A Catalog of Genetic Syndromes in Childhood Cancer. Pediatr Blood Cancer 2015 Dec;62(12):2071–2075.

